# Origins of eukaryotic metabolism

**DOI:** 10.64898/2026.05.08.723234

**Authors:** Carlos Santana-Molina, Anja Spang, Berend Snel

## Abstract

The origin of eukaryotes is a key event in the evolution of cellular life hypothesized to involve a symbiotic integration between a member of the Asgard archaea and the Alphaproteobacteria. Recent work has provided evidence for additional genetic input from other prokaryotes to the eukaryotic proteome yet the extent and sources of these contributions remain debated. Here we aimed to further resolve the prokaryotic origins of eukaryotic genes to inform our understanding of eukaryogenesis. Specifically, we developed a phylogenetic framework to investigate the origins of eukaryotic gene families associated with metabolism and informational processing for comparison. We found that informational processing genes were predominantly derived by archaea whereas eukaryotic metabolism is highly chimeric in its origin. In contrast to previous studies, we report a substantial number of archaeal origins of diverse metabolic enzymes including key metabolic regulators. This highlights an overlooked participation of archaeal metabolism and pinpoints potential metabolic integrations during eukaryogenesis. Apart from the alphaproteobacterial contributions to the eukaryotic metabolism, we found an additional dominant phylogenetic signal of genes potentially derived from Myxococcota, especially for gene families associated with lipid metabolism. By systematically analysing the origins of eukaryotic metabolism, this research offers novel insights into the origin of eukaryotic membranes and refine our current models for the origin of the eukaryotic cell.

## Introduction

The origin of eukaryotes represents a key step in the evolution of the tree of life towards cellular complexity (Szathmáry and Smith 1995). This process, also known as eukaryogenesis, involved a symbiosis between at least two partners: an archaeon from the Asgard archaea (Spang et al. 2015; Zaremba-Niedzwiedzka et al. 2017; Liu et al. 2021; Eme et al. 2023; Zhang et al. 2025), here referred to as Asgardarchaeota (Tamarit et al. 2024) also known as Promethearchaeota (Imachi et al. 2024), and a bacterium from Alphaprotebacteria (Martijn et al. 2018; Muñoz-Gómez et al. 2022) the latter of which was integrated into the cell biology of eukaryotes as the mitochondrion (Roger et al. 2017; Vosseberg et al. 2024). The driving force behind this endosymbiotic integration – syntrophy, phagocytosis or parasitism – is still under debate (Martin and Müller 1998; Cavalier-Smith 2002; Wang and Wu 2014; Spang et al. 2019; López-García and Moreira 2020; Panagiotou et al. 2026), although the main physiological function of mitochondria, among others, suggests that energy production was fundamental for consolidating the symbiosis (Searcy 2003; Roger et al. 2017).

Besides Asgardarchaeota and Alphaproteobacteria contributions, the proteome of the last eukaryotic common ancestor (LECA) has been shaped by contributions from additional prokaryotic groups (Rochette et al. 2014; Pittis and Gabaldón 2016; Méheust et al. 2018; Vosseberg et al. 2021; Bernabeu et al. 2024; Vosseberg et al. 2024; Tobiasson et al. 2025). Indeed, horizontal gene transfer (HGT) or additional transient symbioses are thought to have played a significant role during the emergence of the eukaryotic cell (López-García and Moreira 2020; Donoghue et al. 2023). Based on genomic and *in vivo* evidence, some evolutionary models such as the so called ‘Syntrophy’ (Moreira and López-García 1998; López-García and Moreira 2020) and Entangle–Engulf–Endogenize (E3; (Imachi et al. 2020)) hypothesize a Deltaproteobacteria member (including Myxoccocota) as third participant during eukaryogenesis. In particular, the Syntrophy hypothesis proposes that the Deltaproteobacterium engulfed an archaeal symbiont giving rise to an archaeal-derived nucleus and a bacterial-derived plasma membrane (López-García and Moreira 2020). This would more easily explain the origin of eukaryotic phospholipid cell membranes, which are composed of G3P-linked fatty acids common in bacteria rather than the G1P-linked isoprenoids found in archaea (Lombard et al. 2012). Notably, previous work has indicated that at least a number of eukaryotic genes encoding proteins involved in sterol biosynthesis are derived from Myxococcota (Santana-Molina et al. 2020; Hoshino and Gaucher 2021).

It is generally accepted that genes coding for informational processing machinery, such as replication, transcription, translation, and endomembrane system in eukaryotes have been inherited from the archaeal ancestor of eukaryotes, while genes coding for metabolic proteins are thought to have been predominantly acquired from bacteria, including alphaproteobacteria but also other bacterial groups (Brown and Doolittle 1997; Ribeiro and Golding 1998; Rivera et al. 1998; Jain et al. 1999; Koonin et al. 2004). However, since most evolutionary models of eukaryogenesis propose metabolic interactions between symbiotic partners (López-García and Moreira 2020; Donoghue et al. 2023; Vosseberg et al. 2024), archaeal contributions to eukaryotic metabolism are also expected. In fact, we have recently shown that Asgardarchaeota have made contributions to the central carbon metabolism of eukaryotes: for example some proteins of the glycolytic pathway (acting in the cytoplasm) have archaeal origins, while Alphaproteobacteria have made substantial contributions to the tricarboxyl acid (TCA) cycle (acting in the mitochondria; (Santana-Molina et al. 2025). These observations support the idea that metabolic integration played an important role in eukaryogenesis and that more archaeal contributions to eukaryotic metabolism may yet to be discovered (Santana-Molina et al. 2025; Panagiotou et al. 2026). Identifying these contributions is an overlooked aspect of eukaryogenesis research, but essential for understanding the metabolic core acquired from the archaeal ancestor and the subsequent metabolic remodeling that occurred during the origin of eukaryotic cells.

Here, we use a comprehensive phylogenetic approach to investigate the origins of eukaryotic metabolism in comparison with the origins of informational processing genes for contextualization. Our results show that archaeal contributions to eukaryotic metabolism are sparse, but span amino acid, nucleotide, lipid, and carbohydrate metabolic pathways. In addition, our analyses highlight the participation of additional prokaryotic sources, among which, Myxococcota stood out with a similar number of sisterhood relationships as Alphaproteobacteria and as a major contributor to lipid and amino acid metabolisms. Altogether, this work provides new perspectives focusing on the origins of eukaryotic metabolism which inform current models of eukaryogenesis.

## Results

### Distinct phylogenetic signal for information processing and metabolic genes in eukaryogenesis

We implemented an automatic phylogenetic approach including iterative multiple sequence alignments and a taxonomically representative and balanced database (**Fig. S1**). This approach was used to infer phylogenies for all KEGG gene families assigned to information processing and metabolism. Within each phylogeny, LECA clades were identified using eukaryotic species composition criteria explicitly allowing the presence of interspersed prokaryotic sequences. We inferred the taxonomic composition of LECA sister groups at topological distances of at most 1, 2 or 3, as we noticed many cases indicating that the first sister group may not be sufficient to identify the potential donor clade(s) because of possible phylogenetic artifacts and HGTs (**Fig. S1 and text**). The identification of the sister group was relatively stable when analyzing gene trees using empirical (LG+G) or mixture (LG+C20+G+F) evolutionary models, and showed distinct taxonomic composition for informational processing and metabolic gene phylogenies, respectively (**Fig. 1A,B**).

**Figure 1.**
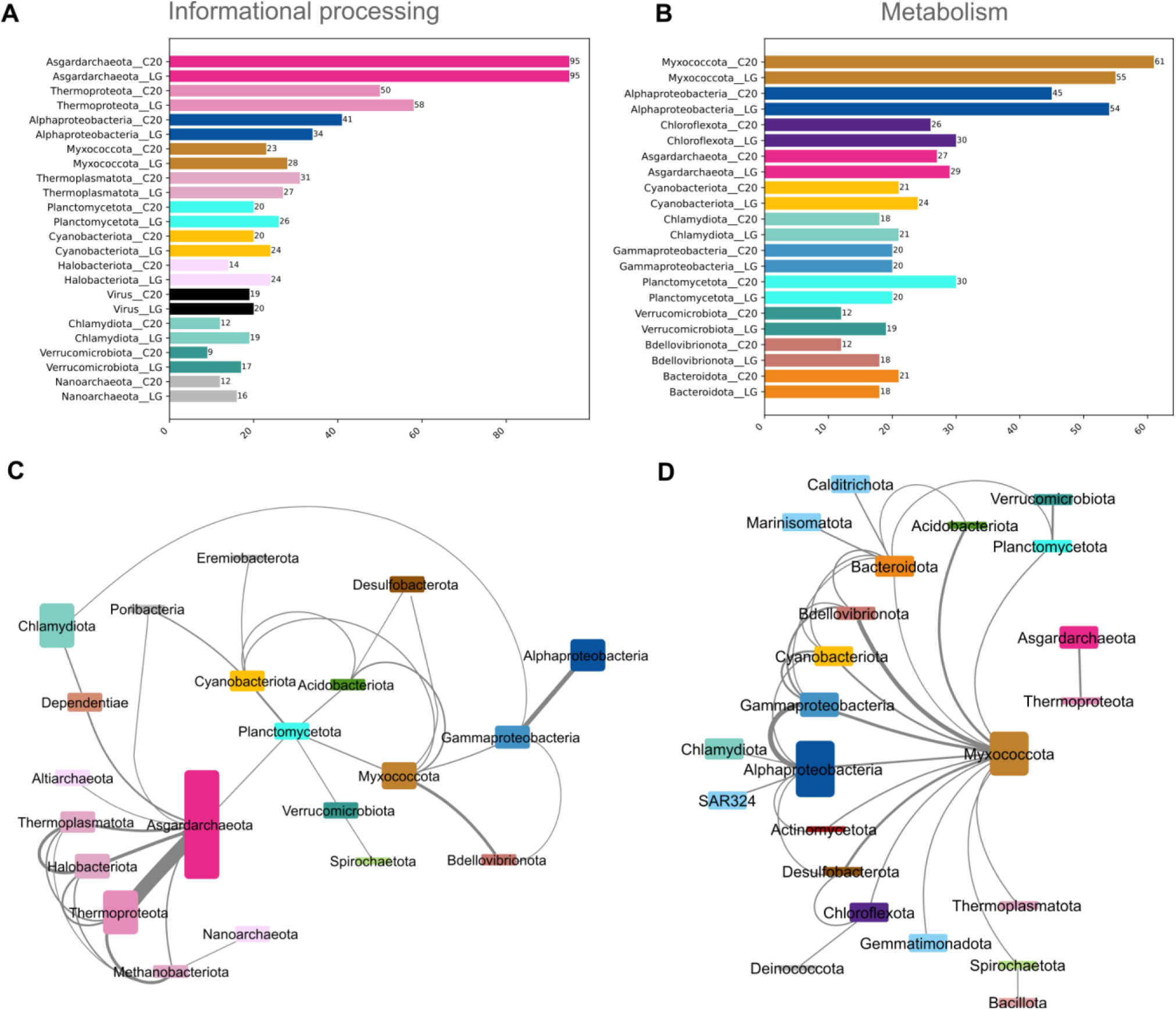
Ranking for the presence and network for the co-occurrences of the most representative prokaryotic phyla in sister groups to LECA clades. Ranking of prokaryotic phyla in sister groups according to phylogenies using LG+C20+G+F and LG+G evolutionary models for **A)** informational-processing and **B)** metabolic related phylogenies. The quantification of presence in sister groups represent an approximation since they are counted for those sisterhood relationships in which the taxonomic proportion is higher than 50%, with ultrafast bootstrap support equal or higher than 95% and for cumulative topological distances 1, 2 or 3. Network for the co-occurrences of prokaryotic phyla in the sister groups to LECA clades for **C)** informational-processing and **D)** metabolic related phylogenies. Co-occurrences were considered at topological distance 3, when the taxonomic proportion of query phylum is higher than 25%. The size of nodes represent the sum of taxonomic proportions for the query phylum while the width of the edges represent the sum of the taxonomic proportion of the target phylum with which co-occur. Only those sum of taxonomic proportions of co-occurrences that sum more than 2 are represented in the network.

Asgardarchaeota are the most prominent phylum among sister groups for informational genes (twice as frequent), followed by Thermoproteota (previously known as TACK; **Fig. 1A**). Alphaproteobacteria form the third most dominant phylum in the sister groups of informational genes, reflecting genes originating from the mitochondrial endosymbiosis such as mitochondrial ribosomal proteins (see below). Given that sister groups are usually mixed taxonomically, we investigated the most common co-occurrences of the prokaryotic phyla in the sister groups to LECA clades at topological distance 3 and represented it in a co-occurrence network (**Fig. 1C and Fig. S2A**). This revealed a clustering by domain with archaeal phyla clustering together but separate from bacterial phyla and vice versa. The clustering of archaeal phyla in the co-occurrence network suggests that eukaryotic clades have sister groups consisting of diverse archaea clades, in which the most common co-occurrence is Asgardarchaeota and Thermoproteota in agreement with vertical inheritance. A similar co-occurrence pattern is observed for Alphaproteobacteria and Gammaproteobacteria. Remarkably, representatives of Chlamydiota and Dependentiae which are not closely related phylogenetically, also often co-occur in the sister groups of LECA clades (**Fig. 1C**), potentially due to shared gene histories as a result of similar lifestyles i.e. parasitic symbionts of eukaryotes (Yeoh et al. 2016; Dharamshi et al. 2023). It is possible that some of these phylogenies represent post-LECA gene exchanges rather than pre-LECA events, but the involved functions are indicative of direct host-symbiont interactions involving for example transcription factors and ubiquitin system related proteins (**Fig. S3**).

Myxococcota and Alphaproteobacteria are the most abundant phyla in LECA sister groups within metabolic gene family trees across varying parameter selection (**Fig. 1B, Fig. S1E**). While the potential contributions from Alphaprotebacteria are expected due to the mitochondrial endosymbiosis, the high number of sisterhood relationships of Myxococcota is notable. We furthermore observed that Cyanoabacteriota, Planctomycetota, and Chlamydiota are present in LECA sister groups in both metabolic and informational protein phylogenies, whereas phyla like Chloroflexota are only represented in LECA sister groups in metabolic gene phylogenies. Asgardarchaeaota are among the top four clades with potential contributions to eukaryotic metabolism highlighting the importance of archaeal metabolism during eukaryogenesis (Santana-Molina et al. 2025). The co-occurrence network of phyla in sister groups of LECA clades in metabolic gene families also reveal a clustering of Asgardarchaeota with Thermoproteota (**Fig. 1D and Fig. S2B**) supporting the vertical inheritance of metabolic genes from the archaeal FECA. However, the co-occurrence of bacterial phyla is more mixed, with Myxococcota being highly connected (**Fig. 1D**), i.e. representing the most common phylum to co-occur with other (bacterial) phyla potentially indicating high degree of HGT involving members of this group. Other bacterial co-occurrences are in agreement with the bacterial species phylogeny, like Myxococcota-Bdellovibrionota, Alpha-Gammaproteobacteria, Planctomycetota-Verrucomicrobiota, Bacteroidota-Marinisomatota consistent with vertical inheritance within bacteria and potential gene transfer to proto-eukaryotes.

### Specific metabolic functions associated with taxa in sister-lineages

We next determined whether any of the phyla found in LECA sister clades are associated with particular metabolic pathways. Specifically, for each major metabolic category, we cataloged the top six potential donor phyla (taxonomic proportion higher than 50% and ultrafast bootstrap higher than 70%) at different topological distances, which revealed excess sisterhood relationships of specific prokaryotic phyla to certain metabolic functions (**Fig. 2A, Fig. S4 and Data S1**). Individual phylogenetic trees corroborate relevant phylogenetic signals as well as the noteworthy information at different topological discances to LECA clades (**Fig. 3** and **4**).

**Figure 2.**
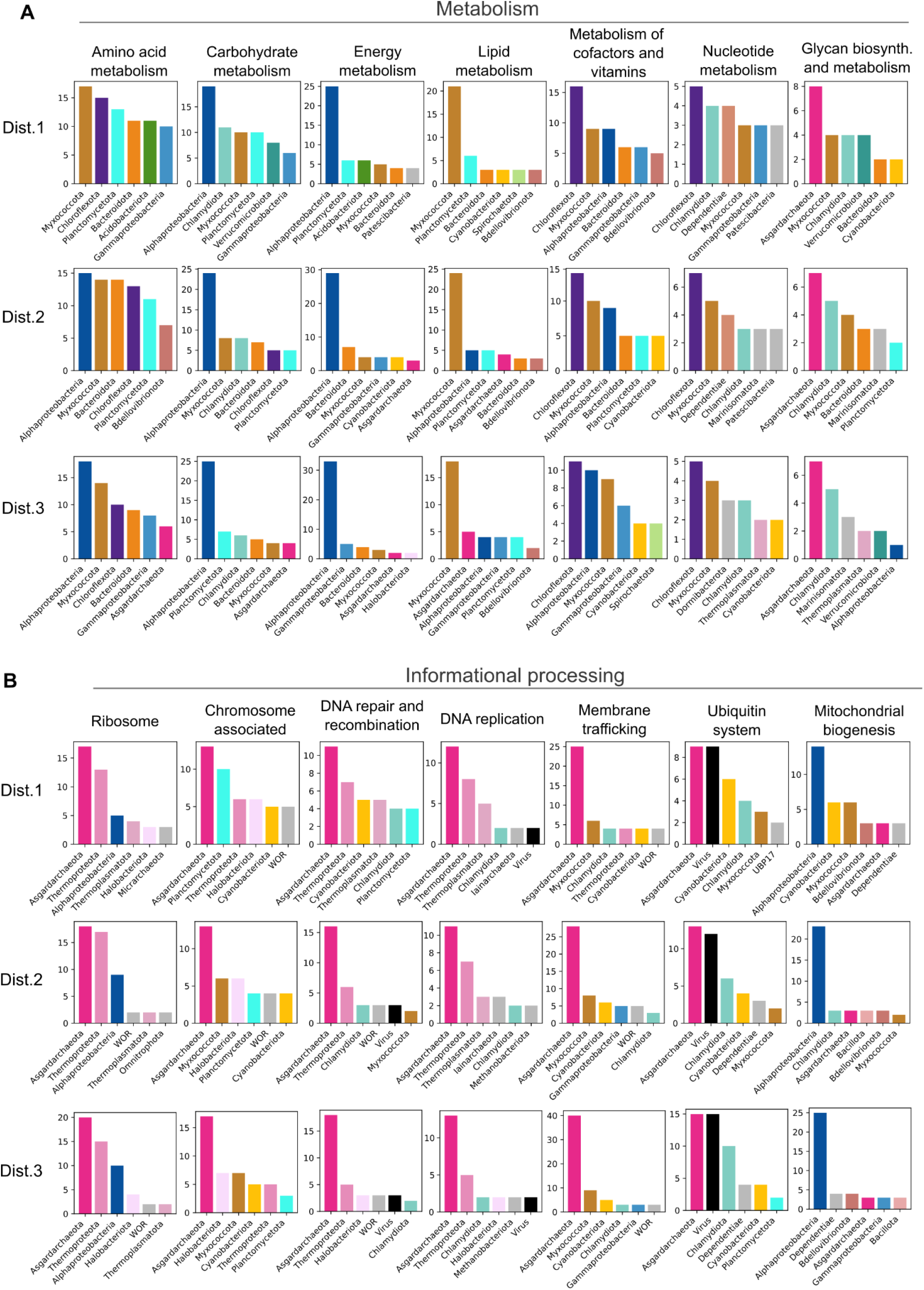
Taxonomic contributions for diverse functional categories related with **A)** metabolism and **B)** informational-processing proteins. Only the top 6 phyla were shown for each category. The sisterhood relationship was considered when taxonomic proportion is higher than 50% and ultrafast bootstrap support equal or higher than 70. Cumulative topological distances 1, 2 and 3, were shown independently. Informational-processing categories were manually selected based on representative functions and a higher number of sisterhood relationships identified. Note that the sisterhood relationship quantification of certain enzymes might be redundant in different categories due to their multifunctional roles.

**Figure 3.**
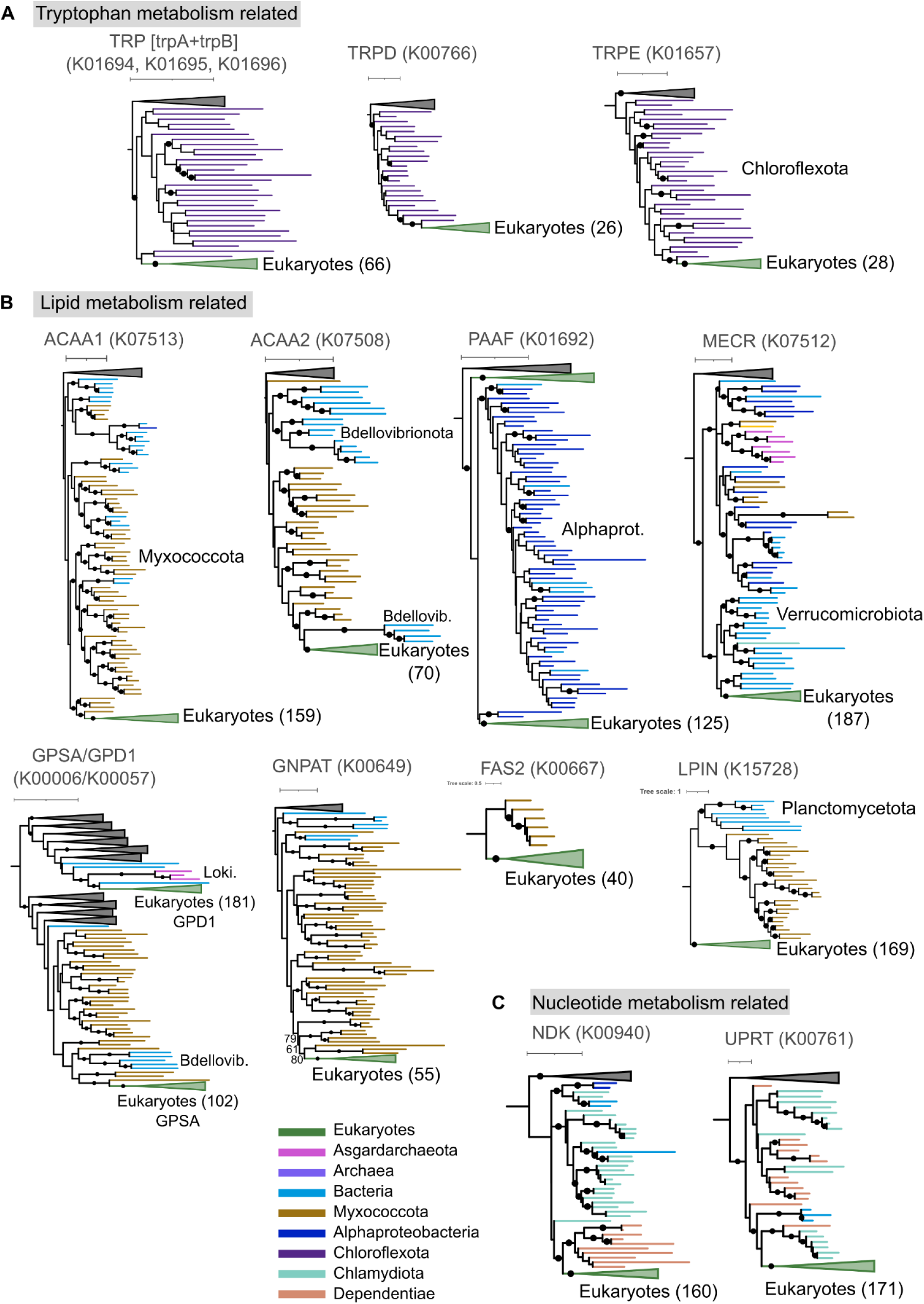
Selected phylogenetic gene trees demonstrating relevant phylogenetic signals of bacterial phyla in sister groups to LECA clades. **A)** Potential gene contribution of Chlorflexota to phenylalanine, tyrosine and tryptophan biosynthesis. TRP tryptophan synthase, result of TrpA and TrpB fusion. **B)** Potential gene contributions of diverse bacterial phyla to lipid metabolism. **C)** Co-occurrence of Chlamydia and Dependentiae in sister groups to LECA clade of nucleotide metabolism related genes. Black dots indicate ultrafast bootstrap higher than 95%. See **Table 1** for gene names and additional information.

**Figure 4.**
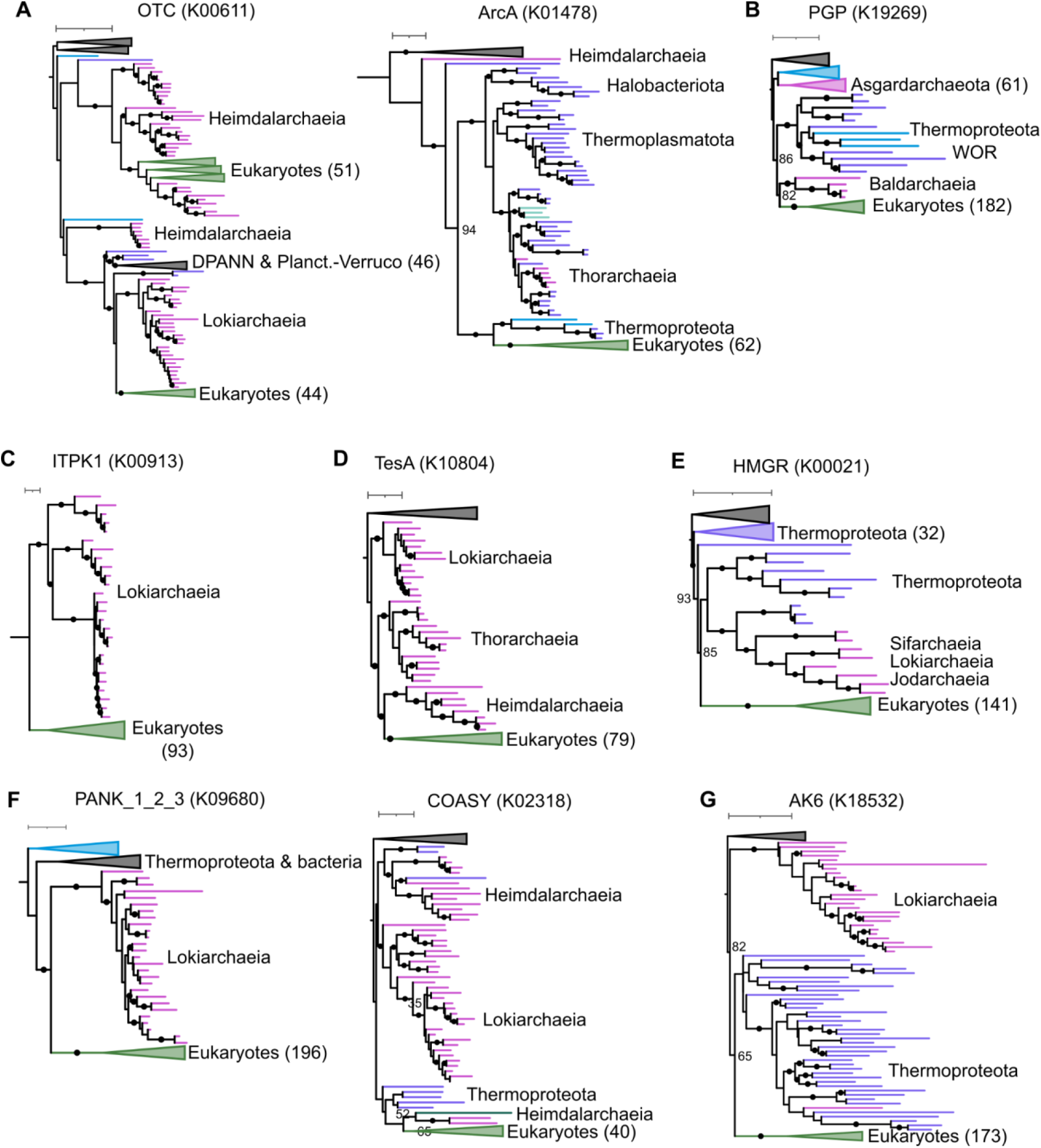
Selected phylogenetic gene trees demonstrating relevant phylogenetic signals of archaea in sister groups to LECA clades including key metabolic regulators. **A)** OTC and ArcA related with arginin metabolism. **B)** ITPK1 regulates inositol metabolism. **C)** PGP regulates glycolate and glycolysis metabolism. **D)** TesA regulates fatty acid metabolism. E) HMGR regulates isoprenoid and sterol metabolisms. **F)** PANK_1_2_3 and COASY regulate pantothenate and CoA biosynthesis. **G)** AK6 regulates ATP, ADP and AMP homeostasis. See **Figs. S5-10** for additional examples of potential archaeal origins in eukaryotic metabolism. See **Table 1** for gene names and additional information.

**Table 1.** Information of relevant enzymes discussed in the text.

| Symbol | KO | Name | Pathways | Sister group |
| --- | --- | --- | --- | --- |
| TRP | K01694 | tryptophan synthase | Phenylalanine, tyrosine and tryptophan biosynthesis | Chloroflexota |
| trpD | K00766 | anthranilate phosphoribosyltransferase | Phenylalanine, tyrosine and tryptophan biosynthesis | Chloroflexota |
| trpE | K01657 | anthranilate synthase component I | Phenylalanine, tyrosine and tryptophan biosynthesis | Chloroflexota |
| trpF | K01817 | phosphoribosylanthranilate isomerase | Phenylalanine, tyrosine and tryptophan biosynthesis | Chloroflexota? |
| TRP1 | K13501 | anthranilate synthase | Phenylalanine, tyrosine and tryptophan biosynthesis | Chloroflexota? |
| OTC | K00611 | ornithine carbamoyltransferase | Arginine biosynthesis | Asgardarchaeota |
| argE | K01438 | acetylornithine deacetylase | Arginine biosynthesis | Asgardarchaeota |
| arcA | K01478 | arginine deiminase | Arginine biosynthesis | Thermoproteota |
| LYS9 | K00293 | saccharopine dehydrogenase (NADP+, L-glutamate forming) | Lysine degradation \| Lysine biosynthesis | Asgardarchaeota |
| paaD | K02612 | ring-1,2-phenylacetyl-CoA epoxidase subunit PaaD | Phenylalanine metabolism | Asgardarchaeota |
| HutI | K01468 | imidazolonepropionase | Histidine metabolism | Asgardarchaeota |
| hisC | K00817 | histidinol-phosphate aminotransferase | Histidine metabolism and others | Asgardarchaeota |
| SEPSECS | K03341 | O-phospho-L-seryl-tRNA<sup>Sec</sup>:L-selenocysteinyl-tRNA synthase | Selenocompound metabolism | Asgardarchaeota |
| ggt | K00681 | glutathione hydrolase | Glutathione metabolism | Asgardarchaeota |
| SDS | K17989 | L-serine/L-threonine ammonia-lyase | Glycine, serine and threonine metabolism and others | Asgardarchaeota |
| TDH | K15789 | threonine 3-dehydrogenase | Glycine, serine and threonine metabolism | Asgardarchaeota |
| CNDP2 | K08660 | cytosolic nonspecific dipeptidase | Arginine and proline metabolism and others | Asgardarchaeota |
| ADPGK | K00874 | ADP-dependent glucokinase | Glycolysis / Gluconeogenesis | Asgardarchaeota |
| GPMI | K15633 | 2,3-bisphosphoglycerate-independent phosphoglycerate mutase | Glycolysis / Gluconeogenesis and others | Asgardarchaeota |
| APGM | K15635 | 2,3-bisphosphoglycerate-independent phosphoglycerate mutase | Glycolysis / Gluconeogenesis and others | Asgardarchaeota |
| ENO | K01689 | enolase | Glycolysis / Gluconeogenesis and others | Aenigmataarchaeota |
| GMPP | K00966 | mannose-1-phosphate guanylyltransferase | Fructose and mannose metabolism and others | Asgardarchaeota |
| PGP | K19269 | phosphoglycolate phosphatase | Glyoxylate and dicarboxylate metabolism | Asgardarchaeota |
| ITPK1 | K00913 | inositol-1,3,4-trisphosphate 5/6-kinase | Inositol phosphate metabolism | Asgardarchaeota |
| ACAA1 | K07513 | acetyl-CoA acyltransferase 1 | Fatty acid degradation and others | Myxococcota |
| ACAA2 | K07508 | acetyl-CoA acyltransferase 2 | Fatty acid degradation and others | Myxococcota |
| HADHA | K07515 | long-chain 3-hydroxyacyl-CoA dehydrogenase | Fatty acid degradation and others | Myxococcota |
| ACADVL | K09479 | very long chain acyl-CoA dehydrogenase | Fatty acid degradation and others | Myxococcota |
| ACADM | K00249 | acyl-CoA dehydrogenase | Fatty acid degradation and others | Myxococcota |
| ACADS | K00248 | butyryl-CoA dehydrogenase | Fatty acid degradation and others | Myxococcota |
| FAS2 | K00667 | fatty acid synthase subunit alpha, fungi type | Fatty acid biosynthesis | Myxococcota |
| ACSBG | K15013 | long-chain-fatty-acid---CoA ligase | Fatty acid biosynthesis and others | Myxococcota |
| PLSC | K00655 | 1-acyl-sn-glycerol-3-phosphate acyltransferase | Glycerolipid metabolism | Myxococcota |
| PLPP1 | K01800 | phospholipid phosphatase 1 | Glycerolipid metabolism | Myxococcota |
| TAZ | K13511 | monolysocardiolipin acyltransferase | Glycerolipid metabolism | Myxococcota |
| GPSA | K00057 | glycerol-3-phosphate dehydrogenase (NAD(P)+) | Glycerolipid metabolism | Myxococcota |
| GNPAT | K00649 | glyceronephosphate O-acyltransferase | Glycerolipid metabolism | Myxococcota |
| SQLE | K00511 | squalene monooxygenase | Steroid biosynthesis | Myxococcota |
| LSS | K01852 | lanosterol synthase | Steroid biosynthesis | Myxococcota |
| CPI1 | K08246 | cycloeucalenol cycloisomerase | Steroid biosynthesis | Myxococcota |
| SMT1 | K00559 | sterol 24-C-methyltransferase | Steroid biosynthesis | Myxococcota |
| CYP51 | K05917 | sterol 14alpha-demethylase | Steroid biosynthesis | Myxococcota |
| LPIN | K15728 | phosphatidate phosphatase LPIN | Glycerolipid metabolism | Myxococcota/Planctomycetota |
| PaaF | K01692 | enoyl-CoA hydratase | Fatty acid degradation and others | Alphaproteobacteria |
| MECR | K07512 | trans-2-enoyl-CoA reductase | Fatty acid metabolism | Verrucomicrobiota |
| PLMT | K00550 | phosphatidyl-N-methylethanolamine N- methyltransferase | Glycerophospholipid metabolism | Asgardarchaeota |
| GPD1 | K00006 | glycerol-3-phosphate dehydrogenase (NAD+) | Glycerophospholipid metabolism | Asgardarchaeota? |
| GBA2 | K17108 | non-lysosomal glucosylceramidase | Sphingolipid metabolism | Asgardarchaeota |
| SGPL1 | K01634 | sphinganine-1-phosphate aldolase | Sphingolipid metabolism | Asgardarchaeota |
| SGPP1 | K04716 | sphingosine-1-phosphate phosphatase 1 | Sphingolipid metabolism | Asgardarchaeota |
| TesA | K10804 | acyl-CoA thioesterase I | Biosynthesis of unsaturated fatty acids | Asgardarchaeota |
| HMGS | K01641 | hydroxymethylglutaryl-CoA synthase | Terpenoid backbone biosynthesis | Asgardarchaeota |
| MVK | K00869 | mevalonate kinase | Terpenoid backbone biosynthesis | Asgardarchaeota |
| MVAK2 | K00938 | phosphomevalonate kinase | Terpenoid backbone biosynthesis | Micrarchaeota |
| MVD | K01597 | diphosphomevalonate decarboxylase | Terpenoid backbone biosynthesis | Micrarchaeota |
| IPK | K06981 | isopentenyl phosphate kinase | Terpenoid backbone biosynthesis | Thermoplasmatota |
| GGPS1 | K00804 | geranylgeranyl diphosphate synthase, type III | Terpenoid backbone biosynthesis | Aenigmataarchaeota |
| DHDDS | K11778 | ditrans, polycis-polyprenyl diphosphate synthase | Terpenoid backbone biosynthesis | Methanobacteriota |
| HMGR | K00021 | hydroxymethylglutaryl-CoA reductase (NADPH) | Terpenoid backbone biosynthesis | Thermoproteota |
| NADC | K00767 | nicotinate-nucleotide pyrophosphorylase | Nicotinate and nicotinamide metabolism | Asgardarchaeota |
| MTHFS | K01934 | 5-formyltetrahydrofolate cyclo-ligase | One carbon pool by folate | Thermoproteota |
| PANK1_2_3 | K09680 | type II pantothenate kinase | Pantothenate and CoA biosynthesis | Asgardarchaeota |
| COASY | K02318 | Coenzyme A synthase | Pantothenate and CoA biosynthesis | Asgardarchaeota |
| rtpR | K00527 | ribonucleoside-triphosphate reductase (thioredoxin) | Purine-Pyrimidine metabolism | Asgardarchaeota? |
| UDP | K00757 | uridine phosphorylase | Pyrimidine metabolism | Thermoproteota |
| AK6 | K18532 | adenylate kinase | Purine metabolism | Thermoproteota |
| UPRT | K00761 | uracil phosphoribosyltransferase | Pyrimidine metabolism | Chlamydiota |
| NDK | K00940 | nucleoside-diphosphate kinase | Purine-Pyrimidine metabolism | Dependentiae |
| IMP | K00088 | IMP dehydrogenase | Purine metabolism | Chlamydiota |

### Amino acid metabolisms

Phylogenies of genes involved in amino acid metabolism show heterogeneous taxonomic sister groups to LECA clades, whose ranking varies depending on the topological distance (**Fig. 2A**). Myxococcota, Alphaproteobacteria, Chloroflexi, Bacteroidota, and Planctomycetota, are most often represented in sister groups of gene family phylogenies associated with amino acid metabolism, and Asgardarchaeota are among the 6 most represented phyla in sister groups when analyzing topological distance 3 (**Fig. 2A**). We found overrepresented taxa for some specific amino acid metabolisms (**Fig. S4A**). For example, we identified 6 potential contributions from Bacteroidota to ‘glycine serine and threonine metabolism’, 7 potential contributions from Chloroflexota to ‘phenylalanine, tyrosine and tryptophan biosynthesis’ (including cooperative enzymes such as TRP [fused trpA/B], trpD and trpE with clear phylogenetic signal (**Fig. 3A**, see **Table 1** for gene names) and trpF and TRP1 with a mixed phylogenetic signal), up to 6 potential contributions from Alphaproteobacteria to ‘tryptophan metabolism’, and 8 possible contributions from Alphaprotebacteria and Myxococcota respectively to ‘Valine, leucine and isoleucine degradation’ (**Fig. S4A**). Furthermore, we found putative contributions to amino acid metabolisms by Asgardarchaeota and other archaeal taxa. These include OTC, arcA (**Fig. 4A**) and argE comprising the final consecutive steps in arginine biosynthesis, LYS9 to lysine metabolism, paaD to phenylalanine metabolism, HutL to histidine metabolism, SEPSECS to selenocompound metabolism, ggt to glutathione metabolism, and other multifunctional enzymes like hisC, SDS, TDH and CNDP2 (**Fig. S5**).

### Carbohydrate and energy metabolism

Alphaproteobacteria appears to be a dominant source of carbohydrate and energy metabolism (**Fig. 2A**) with 30 potential contributions from Alphaproteobacteria mostly associated with oxidative phosphorylation (**Fig. S4B**) and consistent with the crucial role of mitochondria as the powerhouse of eukaryotic cells (Roger et al. 2017). In agreement with previous work (Mahendrarajah et al. 2023), we found that most V-type ATP synthase subunits associated with cellular functions rather than core metabolism are of archeal origin (**Data S1**). We furthermore identified various alphaproteobacterial contributions to pyruvate metabolism and the TCA cycle as reported previously (Santana-Molina et al. 2025). The Planctomycetota-Verrucomicrobiota-Chlamydiota supergroup is frequently observed in sister groups of protein families involved in carbohydrate metabolism (**Fig. S4C**). In addition to previously observed archaeal contributions to glycolysis (e.g. ADPGK, GPMI, APGM and ENO; (Santana-Molina et al. 2025)), we found Asgardarchaeota in sister groups of LECA families involved in nucleotide sugar metabolism (GMPP), glyoxylate metabolism (e.g. PGP (**Fig. 4B**), with an important regulatory and protective role in maintaining the efficiency of glycolysis and other central metabolic pathways) and inositol phosphate metabolism (ITPK1 (**Fig. 4C**), a key regulator in the inositol metabolic pathway; (Desfougères et al. 2019)) (**Fig. S6**). This illustrates the diversity of functions associated with gene families inherited from the archaeal ancestor.

### Lipid metabolism

Myxococcota stands out as a frequent sister lineage of LECA clades mediating enzymatic steps related to eukaryotic lipid metabolism, i.e. we found around 20 potential donations from Myxococcota in this functional category – all other taxa were represented in less than 5 sister clades. These putative Myxococcota contributions to lipid metabolism constitute important enzymatic steps for fatty acid degradation (eg. ACAA1, ACAA2, HADHA, ACADVL, ACADM, ACADS), fatty acid biosynthesis (FAS2, ACSBG), glycerophospholipid metabolism (PLSC, PLPP1, TAZ, GPSA, GNPAT, the two latter are key enzymes in G3P phospholipids linked to fatty acids) as well as sterol metabolism (SQLE, LSS, CPI1, SMT1, CYP51; **Fig. S4D, Data S1**). Myxococcota also seems to represent the only taxon that encodes a full length homolog (∼1,800 amino acids) of FAS2, an enzyme that catalyzes the formation of long-chain fatty acids (**Fig. 3B**) resulting from multiple gene fusions (Bukhari et al. 2014). On the other hand, the phylogeny of other enzymes such as LPIN, showed ambiguous phylogenetic signals with either Planctomycetota or Myxococcota as sister groups, depending on the root (**Fig. 3B**). However not all steps of fatty acid metabolism appear to be of Myxococcota origin. For example, gene families with other taxa observed in sister clades include PaaF with Alphaproteobacteria and MECR with Verrucomicrobiota (**Fig. 3B**). Furthermore, we found up to 6 Asgardarchaeota sisterhood relationships dispersed across lipid metabolic pathways including the biosynthesis of glycerophospholipid (PLMT, GPD1), sphingolipids (GBA2, SGPL1, SGPP1) and unsaturated fatty acids TesA (**Fig. 4D** and **Fig. S7**). This latter enzyme is of particular interest as it plays an important role in regulating the intracellular levels of CoA esters, Coenzyme A, and free fatty acids (Tillander et al. 2017). In addition, we found a substantial number of contributions from archaea specific to the mevalonate isoprenoid pathway: pivotal enzymes like HMGS and MVK shows Asgardarchaea sisterhood relationship, while other enzymes like MVAK2, MVD, IPK, GGPS1, DHDDS and HMGR (the latter a key enzyme that controls sterol biosynthesis in eukaryotes; (**Fig. 4E**)) shows eukaryotic clades nested within archaeal sequences with diverse taxonomy in the sister groups (**Fig. S7**). Together, this indicates that eukaryotic lipid metabolism is of mixed origin similar to central carbon metabolism (Santana-Molina et al. 2025).

### Metabolism of cofactors and vitamins

Chloroflexota seems to be a prominent sister lineage of LECA clades in protein phylogenies related to the metabolism of cofactors and vitamins with up to 16 potential contributions (**Fig. 2A**) acting in nicotinamide, pantothenate, vitamin B6 metabolism and folate biosynthesis among others (**Fig. S4E**). Alphaproteobacteria and Myxococota also have contributed frequently to these functions with Alphaproteobacteria having additionally contributed to ubiquinone biosynthesis (**Fig. S4E**). Furthermore, we found potential asgardarchaeal (and archaeal) contributions to nicotinamide metabolism (NADC), one carbon pool folate (MTFHS), pantothenate and CoA metabolism (PANK1_2_3, COASY; (**Fig. 4F**)), and thiamine metabolism (THI4) (**Fig. S8**). Remarkably, PANK1_2_3 and COASY act in the CoA biosynthetic pathway and are essential for controlling the intracellular CoA concentration as well as the function of mitochondria (Barritt et al. 2024), which illustrate the metabolic integration of archaeal and alphaproteobacterial contributions.

### Nucleotide metabolism

In the nucleotide metabolism, we found a slight overrepresentation of Chloroflexota in sister groups, with 7 and 2 potential contributions to purine and pyrimidine metabolism respectively (**Fig. 2A** and **Fig. S4F**). Further, we identified few archaeal contributions to nucleotide metabolism in phylogenies of RPRT, UDP and AK6 with Asgardarchaeota sequences as sister group, and TMK and PYRG (the later also known as CTP synthetase, a key enzyme in nucleotide metabolism) showing LECA clades nested in archaeal groups (**Fig. S9**). Importantly, AK6 (**Fig. 4G**) catalyzes the interconversion of the various adenosine phosphates (ATP, ADP, and AMP) playing an essential role in cellular energy homeostasis (Ren et al. 2005). We found Chlamydiota in sister groups to LECA clades in phylogenies of key enzymes like IMP dehydrogenase, UPRT and NDK (**Data S1**). The sister groups of the two later gene phylogenies, UPRT and NDK, also included Dependentiae (**Fig. 3C**), which together with Chlamydiota represent well known eukaryotic endosymbionts, suggesting gene exchange of nucleotide metabolism genes with eukaryotes and among these lineages.

### Glycan biosynthesis and metabolism

Asgardarchaeota are a major contributor for glycan biosynthesis and metabolism as we found 4 potential donations to Glycosylphosphatidylinositol (GPI)-anchor biosynthesis, and 7 potential donations to N-glycan biosynthesis (Zaremba-Niedzwiedzka et al. 2017) (**Fig. S10**). Chlamydiota are often in sister groups to glycan biosynthesis and metabolism genes in up to 7 phylogenies - mainly involving Mannose type O-glycan biosynthesis and Glycosaminoglycan biosynthesis (**Data S1**). In addition we found up to 4 potential Myxococcota contributions (**Data S1**), which together with Asgardarchaeota contributions, suggest a chimeric origin of to GPI-anchor biosynthesis involving these two taxa.

### Other metabolisms

Beyond the primary metabolic pathways, we identified distinct phylogenetic signals associated with specific prokaryotic phyla across diverse functional categories (**Fig. S11**). In gene trees related to other amino acid metabolism, Myxococcota and Alpha/Gammaproteobacteria are predominant sister groups to LECA clades. However, the relative ranking of these phyla shifted according to the applied topological distance (**Fig. S11**). Within the biosynthesis of secondary metabolites, Planctomycetota exhibited a slight overrepresentation, appearing as a sister group to LECA in up to six clades. Notably, this phylogenetic signal is lost when the analysis was extended to a topological distance of 3. Conversely, in the metabolism of terpenoids and polyketides, Cyanobacteriota demonstrated a consistent overrepresentation (in up to five clades). This signal remained robust across varying topological distances and subsequent tree inspection suggests that Cyanobacteriota represents a plausible contributor to the LECA proteome. Finally, we observed limited instances of Myxococcota and Asgardarchaeota sisterhood relationships in categories related to membrane transport and catabolism (up to three instances each; **Fig. S11**). It should be noted, however, that the functional versatility of many of these gene families may result in some redundancy with the primary metabolic categories previously discussed (**Data S1**).

### Informational functions associated with taxa in sister-lineages

Asgardarcheota is the most dominant phylum in the sister groups of gene phylogenies associated with many and diverse information processing machinery such as: ribosome, chromosome and associated proteins, DNA repair and recombination, DNA replication, transcription machinery, traslation, membrane trafficking, cytoskeleton and ubiquitin system (**Fig. 2B** and **Fig. S12**). Furthermore, we identified alphaproteobacterial contributions associated with mitochondrial ribosome and biogenesis. We also found up to 3 potential Asgardarchaeota contributions to mitochondrial biogenesis (**Fig. 2B**) including aminoacyl-tRNA biosynthesis genes like YARS, RARS and the endoribonuclease LACTB2, suggesting Asgardarchaeota derived proteins were integrated into mitochondrial cell biology.

Other bacterial contributions are a minority, occur sparsely across informational functions, and inspection of phylogenies show inconclusive sisterhood relationships. We found up to 5 possible contributions from Myxococcota to endomembrane functions (**Fig. 2B**), which might be of relevance in light of the lipid metabolism contributions of this phylum. These contributions include proteins such as SEC13, HPS90A, FAM20B, HDAC6 and SFTPD. However, the phylogenetic signal from these contributions is not always unequivocal, and these proteins belong to different molecular systems. Planctomycetota, was observed in the sister groups of gene phylogenies related to chromosome and associated proteins. However, because this clustering was only observed at topological distance 1 (**Fig. 2B**), we cannot exclude a phylogenetic artifact rather than actual donations from this phylum. WOR-3 phylum was notably present in sister groups of LECA clades in gene trees related with ribosome, chromosome associated proteins, DNA repair and recombination, and membrane trafficking. Inspection of the respective phylogenies suggests that WOR-3 sisterhoods might represent actual donations. Lastly, we found that viruses, Chlamydiota and Dependentiae sisterhood relationship is prevalent in functions related with ubiquitin system which might be related to the fact that members of these are able to hijack the host ubiquitin system to evade the immune response (Viswanathan et al. 2010; Zhou and Zhu 2015).

### Relative timing of acquisition for distinct prokaryotic contributions

To better understand the timing of these various contributions to the emerging eukaryotic cell, we next investigated the normalized stem branch-lengths of LECA clades associated with different donors (Pittis and Gabaldón 2016; Vosseberg et al. 2021). To investigate the stem branch lengths we exclusively relied on sisterhood relationship at topological distance 1 and excluded those LECA clades that contained prokaryotic or viruses sequences interspersed. We subset those sisterhood relationships in which more than 70% of the sister group belonged to the same phyla (considering at least one sequence or more than one sequence, respectively) and have a bootstrap support higher than 70%. Furthermore, we required that LECA clades contain at least 10 (5% of total eukaryotes in the data set) or 60 (25%) eukaryotic representatives, respectively (**Fig. S13**). As reported previously (Vosseberg et al. 2021), we found that stem branch-length of informational genes are usually longer than metabolic gene phylogenies (**Fig. 5**). The stem branch-length of Asgardarchaeota sister groups is generally longer than the stem branch length of other bacterial sister groups - although it is difficult to establish the precise order of acquisition within the bacterial phyla (**Fig. 5A** and **Fig. S13**). This pattern is replicated for gene families encoding proteins associated with informational processing as well as metabolism (**Fig. 5B** and **Fig. S13**). Together, this suggests that Asgardarchaea contributions, including those to metabolism, are the oldest contributions in agreement with the archaeal ancestry of eukaryotes (archaeal FECA; (Pittis and Gabaldón 2016; Vosseberg et al. 2021; Kay et al. 2025)).

**Figure 5.**
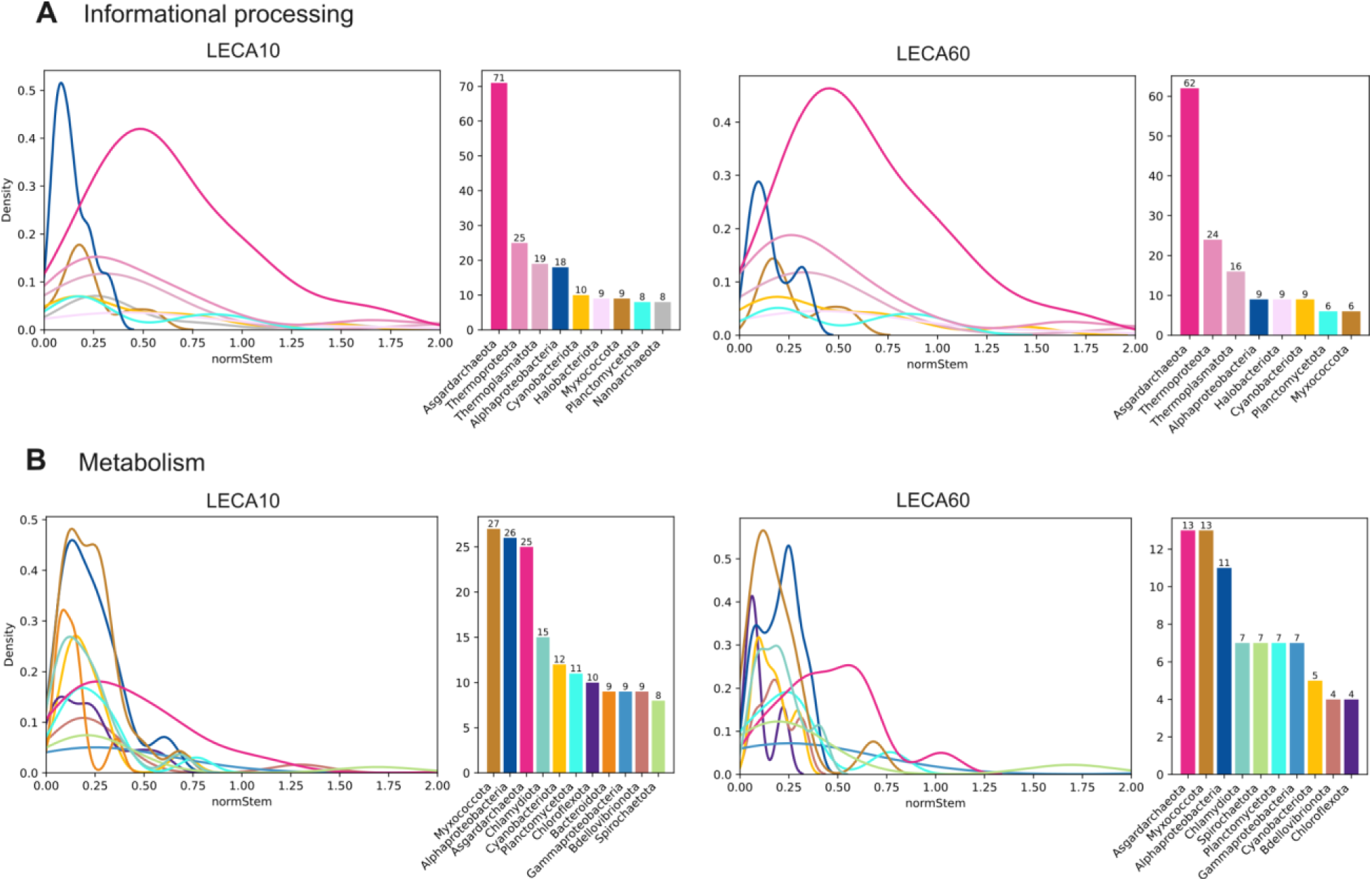
Relative timing of potential gene contributions during eukaryogenesis using normalized stem branch length and different size of LECA clades. Relative timing of **A)** informational processing and **B)** metabolic contributions. Density plots show the distribution of the normalized stem branch lengths of LECA clades and barplots indicate the number of potential contributions representing such distributions. Left panels include those contributions of LECA clades containing at least 10 eukaryotic organisms, and right panels include those contributions to LECA clades containing at least 60 eukaryotic organisms. Only sisterhood relationships at topological distance 1, involving more than 1 sequence, with taxonomic proportion higher than 70% and ultrafast bootstrap than 70%, were analyzed. See **Fig. S13** for extended information.

Our analyses also show that all bacterial taxa have similar ranges of stem branch length when analyzing LECA clades of at least 10 eukaryotes. However, when analyzing LECA clades containing at least 60 eukaryotes, Myxococcota stem branch-lengths appear shorter than those of Alphaprotebacteria (**Fig. 5B**), which would suggest that Myxococcota interaction with proto-eukaryotes postdates Alphaproteobacteria endosymbiosis or that they evolved slower relative to alphaproteobacterial contributions. Planctomycetota contributions appear to involve relatively longer stem branch-lengths than Alphaprotebacteria to LECA clades including at least 60 eukaryotes, but the difference between these estimates are uncertain due to the low number of phylogenies on which they are based (4; **Fig. 5B**). Together, these analyses suggest that Asgardarchaeota (and Thermoproteota) may represent the oldest source of donations to informational processing and also metabolism. Furthermore, although it is difficult to establish a consistent timing of bacterial contributions, we did not find evidence for an early association of Myxococcota and Asgardarchaea preceding alphaproteobacterial endosymbiosis. Instead, our analyses suggest that Myxococcota interaction with proto-eukaryotes may have been concurrent or subsequent to mitochondrial acquisition.

## Discussion

In this work, we implemented a comprehensive approach that analyses sister groups to LECA clades at different topological distances to investigate patterns in the prokaryotic origins of proteins that contributed to informational processing and metabolic functions in the eukaryotic cell. Our analyses reveal that analyzing sister groups at different topological distances we obtain valuable information that provides a consistent and representative view for the diverse phylogenetic signals. In agreement with previous observations (Brown and Doolittle 1997; Ribeiro and Golding 1998; Jain et al. 1999; Koonin et al. 2004; Cotton and McInerney 2010; Méheust et al. 2018), we found that informational processing genes predominantly evolve vertically, which for eukaryotes means inheritance from the archaeal ancestor, while metabolic genes seem to be frequently acquired by endosymbiotic or horizontal gene transfers mainly from bacteria. In addition to these signals consistent with earlier works, our data also revealed new patterns which provide valuable insights into the mosaic origins of eukaryotic proteome and a basis to evaluate current models on the origin of the eukaryotic cell.

### Dispersed metabolic contributions of archaea include key regulators

Combining metabolic and informational processing gene families, we found that Asgardarchaeota is the most dominant phylum with consistent phylogenetic signal in sister groups to LECA clades in agreement with recent analyses (Tobiasson et al. 2026). When considering only metabolic gene families, Asgardarchaeota is still in the top four of sister groups to LECA clades, which is in contrast with the dominant contribution in informational processing gene families. Likewise, we found that the archaeal sister groups encompass diverse phylogenetic signals, such as sister groups consisting of Asgardarchaeaota either next to archaeal clades (eg. PGP, **Fig. 3B**), or next to bacterial clades (eg. GPD1 **Fig. 4B**), as well as presence of additional archaeal groups in sister groups (eg. AK6, **Fig. 3F**). These diverse patterns likely reflect lack of phylogenetic signal in combination with continuous HGT among prokaryotes (Ku et al. 2015) but either way strongly suggest a legacy of contribution of archaeal metabolism during eukaryogenesis. The distribution of the potential origins are dispersed across metabolism including functions such as amino acid, carbohydrate, lipid, nucleotide, cofactor, energy and glycan metabolism. Remarkably some of the retained archeal contributions to the respective metabolisms appear to be metabolic regulators (**Fig. 3**), i.e. enzymes whose activity controls key fluxes through pathways. These key enzymes of potential archaeal origin include PGP for glycolysis (Jeanclos et al. 2022), ITPK1 for inositol (Desfougères et al. 2019), TesA for fatty acid (Tillander et al. 2017), HMGR for sterol (Chen et al. 2022), PANK1/2/3 and COASY for CoA biosynthesis (Barritt et al. 2024), as well as AK6 for ATP/ADP/AMP homeostasis (Ren et al. 2005). This pattern suggests that the inheritance of metabolic regulators may have been key for the subsequent metabolic integrations during eukaryogenesis as they might have played a role in the functional control of enzymes derived from the bacterial contributions. On the other hand, we did not find archaeal contributions related with eukaryotic aerobic metabolisms such as CoXABC and Qcr2 (Appler et al. 2026), which instead, have been acquired from the Alphaproteobacteria. The oldest timing of the asgardarchaeota origins to metabolism, as suggested by their stem branch-lengths, together with their incidental presence in diverse metabolic pathways is strongly suggestive of an archaeal-FECA metabolism that over time integrated various bacterial contributions and was shaped by gene loss and replacement.

### The origin of eukaryotic cell membranes is related with Myxococcota gene contributions

The abundance of Myxococcota in the sister groups to metabolic LECA clades is noteworthy and comparable with the number of Alphaprotebacteria contributions (**Fig. 1C**). Our results not only suggest abundant potential donations from Myxococcota but also the involvement of members of this group in frequent HGT with diverse other bacterial phyla in the sister groups to LECA clades (**Fig. 1D**). These abundant Myxococcota contributions may have been overlooked due to the low representation of members of this taxon in previous analyses. Notably, our analyses suggest that Myxococcota have made significant contributions to lipid metabolism involving degradation of fatty acids, biosynthesis of phospholipids of fatty acids and sterol metabolism (the latter in agreement with previous analyses (Santana-Molina et al. 2020; Hoshino and Gaucher 2021)). As we found an archaeal origin of the eukaryotic mevalonate isoprenoid pathway (Zhu et al. 2024; Tobiasson et al. 2025), the sterol biosynthesis acquired from Myxococcota likely used isoprenoid precursors derived from the metabolism of the archaeal ancestor of eukaryotes which illustrate metabolic integrations between myxococcota and archaeal donors. These observations related with lipid metabolism, especially the finding that eukaryotic phospholipid biosynthetic genes such as FAS2, GPSA and GNPAT may derive from Myxococcota provide critical new data to address the highly debated lipid membrane transition during eukaryogenesis (Lombard et al. 2012). In addition, it is notable that at least 8 potential contributions are related to fatty acid degradation. This pathway in eukaryotes represents an alternative for fuelling carbon and energy metabolism (fatty acid beta oxidation) which might have been metabolically favorable for early eukaryotic cells. Myxocooccota contributions at first glance seem to align with the Syntrophy hypothesis (Moreira and López-García 1998; López-García and Moreira 2020). However, we did not identify strong evidence for contributions of Myxococcota to eukaryotic information processing machinery suggesting that Myxococcota contributions specifically provide metabolic functions. Furthermore, while the Syntrophy hypothesis posits that the Myxococcota interaction with Asgardarchaeota predates the mitochondrial endosymbiosis, our results on the stem branch-lengths suggests that this interaction might have taken place concurrent or after mitochondrial integration. This timing is consistent with recent work that points to an autogenous development of at least some of defining features of complex eukaryotic cells (including the building blocks of a nucleus) within the Asgardarchaeal ancestor of eukaryotes prior to mitochondrial acquisition (Vosseberg et al. 2021; Kay et al. 2025) and some of the cell biological features of extant Asgardarchaea (Imachi et al. 2020; Rodrigues-Oliveira et al. 2023; Radler et al. 2025). These observations and other recent work are more easily reconcilable with the inside-out hypothesis in which the nucleus evolved from the cell body (Baum and Baum 2014; Baum and Spang 2023). Thus, Myxoccocota may have been additional symbiotic partners of the emerging eukaryotic cell, which happened to contribute lipid metabolism genes to the Asgardarchaea-derived nuclear genome that eventually led to the replacement of archaeal lipids in both the cell membrane and intracellular organelles. However, because genes evolve at different rates in different organisms and our analyses did not integrate molecular dating approaches, further work is needed to establish an absolute timeline for Myxococcotal contributions compared to those of Alphaproteobacteria and further discriminate between these two and other models on eukaryogenesis.

### Reciprocal gene exchange of Chlamydia and Dependentiae parasites with eukaryotes

Our results furthermore provided new evidence for gene exchange between Chlamydiota and Depedentiae parasites in early eukaryotic evolution. While Chlamydiota has previously been shown to have exchanged genes with eukaryotes (Becker et al. 2008; Stairs et al. 2020), the gene exchange between Depedentiae and eukaryotes has not been noted so far. We found that Chlamydiota and Dependedentiae often co-occur in the sister group of LECA clades of informational processing genes involving transcription factors, tRNA synthases, and ubiquitin related functions. The presence of ubiquitin components in intracellular parasites (and viruses) is known as a strategy to hijack the host ubiquitin system in order to enhance replication, evade the immune response, and increase pathogenesis (Zhou and Zhu 2015). Given that ubiquitin system is a signaling system exploited in eukaryotes (and also found in Asgardarchaeota **Fig. 2B**) and taking into account the evolutionary history of respective genes (**Fig. S3**), the most plausible explanation is these prokaryotes (and viruses) acquired such genes from (proto-)eukaryotes, rather than prokaryotes or virus HGT to eukaryotes. On the other hand, from a metabolic perspective, these two phyla also co-occur in phylogenies of key enzymes for nucleotide metabolism such as UPTR and NDK (**Fig. 4C**). In addition, we found a number of exclusive sisterhood relationships of Chlamydiota in functions associated with carbohydrate, nucleotide and glycan metabolisms with phylogenies strongly suggesting HGT from Chlamydiota to LECA. Thus, the phylogenies containing these sister groups do not allow for a single interpretation, i.e it could be possible that Chlamydia and Dependentiae exchanged genes and later transferred genes to proto-eukaryotes, that Chlamydia and/or Dependentiae acquired the genes from proto-eukaryotes or a combination of both options i.e. reciprocal gene exchange. Regardless of the direction of the transfers, since Chlamydiota and Dependentiae are unrelated taxa in the bacterial tree of life, the co-occurence of both in sister groups to LECA clades suggests a convergent acquisition of eukaryotic genes warranting further exploration. Furthermore, these cases illustrate that intracellular parasitism is a potential source of gene exchange during and after eukaryogenesis.

### Eukaryotic metabolisms are highly chimeric in their origins

The phylogenetic trees contain strong evidence for substantial contributions from many other prokaryotic taxa to specific metabolic functions. Notably, members of the phylum Chloroflexota seem to represent important players in the provision of genes associated with the metabolism of cofactors/vitamins, nucleotides, and specific amino acids, including phenylalanine, tryptophan, tyrosine, arginine, and lysine. In phylogenetic analyses of multiple pathways for carbohydrate, lipid, and amino acid metabolism, the Planctomycetota and Verrucomicrobiota phyla consistently appear in sister groups. Although the phylogenies of amino acid metabolism reveals heterogenous origins, we found substantial contributions of Bacteroidota, especially to glycine, serine, and threonine metabolisms. However, while many of these contributions could be genuine and represent early HGTs to the emerging eukaryotic cell from various sources, we cannot rule out phylogenetic artifacts, ongoing HGT within prokaryotes and between prokaryotes and early eukaryotic cells, or lack of signal obscuring the identification of real donors.

Contributions such as the tryptophan biosynthesis genes are potentially indicative of a single event which, if not endosymbiotic gene transfer, could be operon transfer. Validation of a single event would allow concatenation of the individual enzymes which would provide more accurate sister group identification and better branch length timing. Likewise, although many of the phyla we found in sister groups align with recent studies on the origin of the eukaryotic proteome (Vosseberg et al. 2021; Bernabeu et al. 2024; Tobiasson et al. 2026), we did not observe the same overrepresentation of some groups like Actinomycetota (Tobiasson et al. 2026). This suggests that findings in this field may be heavily influenced by the specific datasets and methods used, and thus, these contributions require further detailed investigation.

## Conclusions

Attributing the evolutionary origins of individual enzymes in eukaryotic proteomes has been largely explored through a lens in which Alphaproteobacteria and Asgardarchaeaota are the main players involved. However, the increasing availability of extensive prokaryotic diversity is allowing us to develop a more detailed and biologically accurate model of eukaryogenesis. In this work, we focus on the origin of eukaryotic metabolisms to deepen our understanding of the early evolution of the eukaryotic cell and potential interactions of Asgardarchaea with other prokaryotes that have shaped subsequent metabolic integrations. Our results reveal that the origin of eukaryotic metabolism is highly chimeric, and identify Asgardarchaeaota origins as the potentially oldest metabolic substratum. It seems likely that the original asgardarchaeal metabolism was slowly replaced and complemented by bacterial contributions, especially after the acquisition of the alphaproteobacterial endosymbiont forming the mitochondrion. Genes that were retained from this archaeal ancestor include among others, key metabolic regulators that may have ensured the subsequent metabolic integration and physiological control. Furthermore, we show that Myxococcota may represent additional symbiotic partners whose contributions have shaped the origin of eukaryotic cell membranes. In turn, our catalog of phylogenetic trees contextualizes eukaryogenesis as a multi partner affair with diverse prokaryotic taxa contributing to the origin of eukaryotic metabolic pathways.

## Methods

### Datasets constructions

#### Proteome selection and processing

We assembled protein families by using two datasets for different purposes which are i) core_datasets for the identification of LECA clades, and ii) expanded_dataset for the identification of sister groups. First, we employed an updated version of the previous dataset (Santana-Molina et al. 2025) increasing the Asgardarchaea representatives, and comprising a total of 661 archaea, 487 bacteria (19 archaeal and 95 bacterial phyla) and 224 eukaryotic proteomes downloaded from EukProt v3 (Richter et al. 2022). Note that these eukaryotic proteomes were filtered out for redundant sequences as well as for eukaryotic contaminations (Santana-Molina et al. 2025). In addition, we filtered out potential prokaryotic contaminations from eukaryotic proteomes by doing DIAMOND v.2.0.6 (Buchfink et al. 2021) protein searches of eukaryotic proteomes against the NCBI_nr database (release 244). For each sequence we analyzed the top hits: if the first top hit is prokaryotic and has an identity > 0.8 and the alignment cover is > 0.45, the sequence was considered as prokaryotic contamination; likewise, when these thresholds failed and 80% of the top 10 hits was prokaryotic, the sequence was considered as contamination as well. This dataset was denoted as *core_dataset*, it is taxonomically unbalanced (**Fig S1A**) and was exclusively used for the identification of major LECA clades in phylogenetic trees using KO gene families (see below). The second dataset was constructed by the selection of representative proteomes of GTDB (release 207; (Parks et al. 2022)) and viruses proteomes from NCBI (release 244), to which we refer as *expanded_dataset* (**Fig S1A**). For the bacterial selection of this *exapanded_dataset*, we sampled proteomes at order level: if the number of taxonomic orders within a phylum is higher than 100, we selected one proteome per order with the higher completeness, the lower contamination, and type-strain if possible (those with genome completeness lower than 80% and contamination higher than 5% were excluded, excepting for Patescibacteria for which we consider those genomes with higher completeness than 60%). For those phyla in which the number of orders is lower than 100, we sampled at genus level, by selecting the top quality genomes with the same threshold as above. If the sampled genomes for a phylum was still lower than 50 we made a second round of sampling. The rationale of this iterative sampling was to achieve a more taxonomically balanced and complete dataset. For the selection of archaeal proteomes, due to genome reduction being more prominent specially in DPANN organisms, we selected genomes by sequence identity of the concatenated gene markers provided by GTDB and which exclusively contains representative taxa. We clustered the concatenation by taxonomic classes, and reduced redundancy of the concatenation at sequence identity of 85% using trimAL (-maxidentity 0.85; (Capella-Gutiérrez et al. 2009)), using the remaining proteomes as representative for our final dataset. In total, the expanded_dataset consisted of 154 bacterial phyla (3,520 proteomes), 20 archaeal phyla (1,394 proteomes), 44,289 viral proteomes, plus the 224 eukaryotic proteomes previously selected.

### KO annotation and classification

Core_dataset was annotated using KOFAM_SCAN v.1.3.0 (-f mapper-one-line, e value 1e−3; (Aramaki et al. 2020)) and HMMER.3.2.3 (e-value 1e−3, selecting best i-value hit); (Potter et al. 2018)), against KO.hmm database. To identify sequences for metabolic gene trees, we primarily used Kegg orthology (KO; Supplementary Data 2) annotations, prioritizing KOFAM classifications.

In instances where KOFAM annotation was absent, we relied on HMMSEARCH annotations. For the distinction of Informational *versus* Metabolic genes, parsed the KEGG hierarchy file (ko00001.keg). Informational gene families were those belonging to ‘Genetic Information Processing’ hierarchy, and those belonging to ‘Brite Hierarchies’ with the following categories: Transcription factors, Transcription machinery, Messenger RNA biogenesis, Spliceosome, Ribosome, Ribosome biogenesis, Transfer RNA biogenesis, Translation factors, Chaperones and folding catalysts, Membrane trafficking, Ubiquitin system, Proteasome, DNA replication proteins, Chromosome and associated proteins, DNA repair and recombination proteins, Mitochondrial biogenesis. Metabolic gene families were those belonging to the ‘Metabolism’ hierarchy. Only those KO sets that contain at least 10 eukaryotic sequences were selected for downstream analyses. In total, we analyzed 4,606 informational processing, and 3,500 metabolic KOs.

### Merging Kos

Because KO classification in many cases is based in functional rather than in evolutionary relationships, we combined KO sets based on HMM homologies using data generated in previous analysis (Santana-Molina et al. 2025). We systematically merged KOs when a KO set (core_dataset) predominantly contained eukaryotic sequences (>90%) and HHSEARCH search (HH-suite 3.1.0 (Steinegger et al. 2019)) indicated a high probability (>90%) and coverage (> 80%). For each query KO, we clustered the two closest target KOs, and merged those KO sets by homology network connectivity using networkx python library. This clustering resulted in 3,920 Informational and 2,868 metabolic KO clusters.

### Phylogenetic reconstructions

We performed a series of iterative multiple sequence alignments (MSAs) in order to get accurate phylogenetic reconstructions. We first used the clustered KO sets from the core_dataset for making phylogenies and to identify the 4 biggest LECA clades (see below), and then used the expanded_dataset for making protein sequence searches and phylogenies oriented towards the sister groups of the selected LECA clades (**Fig. S1B**).

### First round with core_dataset

MSAs were constructed iteratively in order to align sequences with an accurate method and reduce computational time. We first used MAFFT -auto (Katoh and Standley 2013) combined with a strict trimming using trimAL (first we applied a gap trimming with -gt 0.65 option, and then we removed unconserved positions and sequences with -resoverlap 0.70 -seqoverlap 80 options). The respective MSAs were used for making HMM using HMMBUILD (Potter et al. 2018), to which sequences were realigned using HMMALIGN (-trimm). The resulting alignment in stockholm format was converted into fasta unaligned sequences removing any sequences shorter than 80% of the average length of the original KO set. Then, we realigned sequences MAFFT-LINSI (Katoh and Standley 2013), trimmed the gaps with trimAL (-gt 0.7), removed redundant sequences with trimAL (-maxidentity 0.90), and removed further positions with low conservation using trimAL (-automated1). This process was unsuccessful for some KO sets retaining very few positions, and thus we used an alternative method. Those final alignments that retained less positions than 30% of the average length of full KO set, or retained less than 30% of sequences after the trimming, were realigned using MAFFT -auto, trimmed the gaps using trimAL (-gt 0.40), and then removing unconserved positions and spurious sequences using trimAL (-resoverlap 0.50 -seqoverlap 50). The respective alignments were used for phylogenetic reconstructions using IQ-TREE 2.1.2 (Minh et al. 2020) using LG+G model and ultrafast bootstrap (-B 1000).

### Second round with expanded_dataset

Using a custom ETE4 (Huerta-Cepas et al. 2016) script, we identified the potential LECA clades (see below) across the phylogenies generated with the core_dataset. In total, we analyzed 1,830 informational and 1,430 metabolic clustered KO sets that contained at least one LECA clade. We selected the 4 top biggest LECA clades for each KO set, and constructed eukaryote-specific HMMs by aligning with MAFFT -auto (Katoh and Standley 2013), trimming gaps with trimAL (-gt 0.4), and using HMMBUILD (Potter et al. 2018). We performed HMMSEARCHES with an e-value threshold of 1e-5 against the prokaryote specific expanded_dataset, extracting the top 200 sequences sorted by i-value per each search. The target sequences plus the sequences forming the LECA clades belonging to the same clustered KO set, were merged for final phylogenetic reconstructions and MSAs were conducted iteratively as described above. Final phylogenies were performed with IQ-TREE 2.1.2 (Minh et al. 2020) using LG+G, and LG+C20+G+F independently, and using ultrafast bootstrap (-B 1000).

### Identification of sister groups

We systematically analyze phylogenetic trees with a custom ETE4 (Huerta-Cepas et al. 2016) script to identify the potential LECA groups, the respective sister groups at different topological distances i.e number of nodes regarding one specific node, as well as different parameters associated to them like the bootstrap support values, taxonomic composition, branch-lengths, and composition of LECA clades.

### Gene tree rooting

For each individual phylogenetic tree, we performed two rooting steps. Firstly we root the tree at the largest prokaryotic monophyletic group to identify the potential LECA clades (see below). Secondly, for each potential LECA clade, we root the most distant leaf to the common ancestor node of the LECA clade and do not belong to the respective LECA clade.

### LECA clade identification

Isolated prokaryotic sequences can branch within potential LECA clades, provoking the fragmentation of these into multiple monophyletic eukaryotic clades that belong to the same LECA clade. To tackle these cases, we clustered those eukaryotic monophyletic groups which are at topological distance 2 by network connectivity. Thus, the LECA clades that we identify might be composed of monophyletic and/or paraphyletic groups of eukaryotic sequences. Then, we defined the LECA clades as phylogenetic clusters of eukaryotic sequences that include at least 10 organisms representing at least two of the major eukaryotic groups: Excavata, Amorphea, and Diaphoretickes. Clusters that exclusively contained Diaphoretickes and Euglenids were excluded as they potentially represent secondary endosymbiont acquisitions. For the functional annotation, we chose the most abundant KO annotation among the eukaryotic sequences forming the respective LECA clade.

### Sister groups at different topological distances

For each LECA node with its respective most distant rooting, we analyzed the prokaryotic/virus sister groups that were at topological distance 1, 2 and 3 in a cumulative fashion. Thus, we analyzed the sister groups at distance, 1, 1-2 and 1-2-3. From these distance levels we extracted information like taxonomic proportion of phyla i.e. number of sequences belonging to a specific phylum out of the total of sequences forming the sister group at such distance; presence of taxonomic classes and orders per each phylum; the minimum distance bootstrap support per phylum i.e. the support of the closest distance for a specific phylum; the maximum bootstrap support i.e. the maximum support of a specific phylum at any of the distances (1, 2 or 3); proportion of bacterial versus archaeal sequences forming the sister group(s); prokaryotic/virus sequences branching within potential LECA clades; as well as information associated to eukaryotes like size of the LECA clade, and stem branch-length of LECA clade.

### Normalized branch-length analyses

To investigate the relative timing of the potential contributions, we used normalized stem branch-length of LECA clades following the previous approach (Pittis and Gabaldón 2016), that is the relation between stem branch-length of the LECA clades and the median of the branch lengths within the LECA family (from tips to LECA node). We exclusively analyzed the sisterhood relationships at topological distance 1 (taxonomic proportion > 70% and ultrafast bootstrap > 70%), and those LECA clades that did not contain interspersed prokaryotic sequences.

## Author contributions

This study was conceived by C.S-M., A.S., B.S. Research was supervised by A.S and B.S. Experiments and data analyses were carried out by C.S-M. The manuscript was written by C.S-M., A.S., B.S. All authors read and approved the final manuscript.

## Acknowledgments

We thank collaboration partners of this project, E. J. Javaux, P. Mason and R. Hennekam.

## Code and data availability

Phylogenetic trees, code and data of the analyses is available at zenodo repository (https://doi.org/10.5281/zenodo.19234485).

