## Supplementary material for "Origins of eukaryotic metabolism": Fig. S

#### Identification of sister groups to LECA clades considering different parameters

To investigate the origins of eukaryotic metabolism, we used functional annotations denoted by KEGG database, and selected those gene families belonging to *Metabolism* hierarchy as well those belonging to *Informational processing* hierarchies (*see methods*) to contrast the different phylogenetic signals. We iteratively refined our multiple sequence alignments (MSAs) and utilized a taxonomically balanced database to generate phylogenetic trees with a robust phylogenetic signal and reliable sister group representation. We first used a small representative dataset (*core_dataset;* **Fig. S1A** left panel) to construct the phylogenies of the gene families and identify the potential LECA clades. For each of these KO set phylogenies, we selected the 4 biggest LECA clades (defined by phylogenetic clusters of eukaryotic sequences that include at least 10 organisms representing at least two of the major eukaryotic groups: Excavata, Amorphea, and Diaphoretickes; *see methods*), built HMMs and performed HMMSEARCHES against an expanded and taxonomically balanced database (expanded_dataset, *see methods*; **Fig. S1A** right panel) to reconstruct phylogenies oriented towards LECA clades with the idea of improving the phylogenetic resolution of sister groups (Methods; **Fig. 1B**).

Investigating the evolution of metabolic genes is challenging because they are more prone to horizontal gene transfers (HGTs) than informational processing related genes (*see main text*). Particularly, the evolutionary history of genes more prone to HGT including during eukaryogenesis might result in phylogenetic trees with ambiguous sisterhood relationships interpretations (**Fig. S1C**). The phylogeny of a gene that has been transferred from Asgardarchaeota to other prokaryotes and ending up in proto-eukaryotic (pre-LECA) lineages, can provide similar topologies as a gene than has been vertically transferred from Asgardarchaeota to proto-eukaryotes with subsequent HGTs from proto-eukaryotes to prokaryotes (**Fig. S1C**). To aid in the interpretations of these confounding scenarios for gene contributions to LECA, we have analyzed the taxonomic composition of the sister groups to LECA clades that are at topological distances 1, 2 and 3, in a cumulative fashion (**Fig. S1D**). Thus, for each of these distances and for each present phylum, we stored information like taxonomic proportion, minimum distance bootstrap support, bootstrap support and features associated to eukaryotes like size of LECA clades as well as stem branch-lengths (**Fig. S1D**).In addition, our identification of LECA clades consist of the clustering of eukaryotic monophyletic groups that are at topological distance 2, which allow the detection of more representative LECA clades due to we often observed prokaryotic sequences interspersed (*see methods*).

Using this approach we were able to monitor the taxonomic compositions in sister groups comprehensively. Specifically, we combined cumulative topological distance (1, 1-2, 1-2-3), bootstrap support, and taxonomic proportion to plot the presence of most abundant phyla in sister groups to LECA clades for genes involved in informational processing  and metabolism, respectively (**Fig 1E,F**). This representation shows a substantial decrease of the presence of phyla when considering bootstrap thresholds (0, 70, 95%), and when combining it with different taxonomic proportion thresholds (25, 50 and 90%). Although this representation shows the presence of phyla, we might consider those presences with >70% UfBoot2 and >50-70% of taxonomic proportion as tentative donations (black asterisks in Fig. 1D) and those with >95% UfBoot2 and >90% of taxonomic proportion as likely donations (gray asterisks in **Fig. 1E**). For the informational processing set of genes, the comparison of these thresholds at different distances shows an increase for the detection of tentative donations when analyzing the sister groups at distance 3 compared to lower distances (horizontal discontinuous line in **Fig. 1E**), which illustrate the potential use of this approach. By contrast, the detection of likely donations remains relatively stable across the distances (horizontal discontinuous line in **Fig. 1E**). When comparing the presence of phyla for metabolic and informational processing sets of genes, we observe a clear distinction in the taxonomic compositions of the sister groups to LECA clades (**Fig. 1D,E**). Informational processing protein phylogenies clearly shows Asgardarchaeaota as most dominant phylum in the sister groups while metabolic protein phylogenies shows a more heterogeneous compositions in which Myxoccocota and Alphaproteobacteria are the ones slightly more prevalent. In addition, for the metabolic set of genes, we found that phyla like Planctomycetota are enriched in sister groups, but their presence drastically decreases when considering taxonomic proportion metrics. This suggests that this phylum is often present in sister groups but in co-occurrence with other taxonomic groups indicative of HGT among prokaryotes.

**Supplementary Tables**

**Data S1.** Data for the description of sisterhood relationships of LECA clades for metabolism and informational-processing related phylogenies. Each row correspond to a specific LECA clade, with the following information: LECA_size denotes number of eukaryotes forming the LECA clade, NormStem denotes the normalized stem branch-length, D1/D2/D3_phylum denotes the most abundant phylum in the sister groups at different topological distance, _Support denote the ultrafast bootstrap support for the sisterhood relationship, _txporp denotes the proportion of the most abundant phylum in the sister groups, _ntax denotes number of sequences for the respective taxonomic proportion, ArchaeaPropD3 and BacteriaPropD3, denotes the proportion of archaeal and bacterial sequences at topological distance 3. 'na' values indicate sisterhood relationships that contain eukaryotic sequences at the given topological distance. Note that phylogenies containing more than 15 LECA clades were excluded since they most likely represent complex phylogenetic relationships.

**Supplementary Figure Legends**

**Figure S1.** Dataset compositions, methodology and validation. **A)** Taxonomic composition of the datasets employed, core_dataset and expanded_dataset (*see methods*). **B)** Methodology to obtain phylogenetic trees based on two main steps. First step uses core_dataset for detecting LECA clades and second step uses expanded_dataset to obtain representative sister groups of LECA clades. **C)** Two evolutionary scenarios of gene flow that can result in the same tree topology. **D)** Schematic phylogenetic tree to show the parameters to consider in sister groups to LECA clades at different topological distances. Monitoring the presence of prokaryotic phyla in sister groups to LECA clades and validation of analyzing sister groups at different topological distances for **F)** informational and **G)** metabolic gene trees. Each data point represents different parameters considered, where X labels contain information for the cumulative distance (1, 2, 3), ultrafast bootstrap (bs70, bs95), and taxonomic proportion (txp25, txp50, txp90). Discontinuous horizontal lines and asterisks show the increasing detection of phyla in sister groups at different topological distances and with the same taxonomic proportion and ultrafast bootstrap parameters.

**Figure S2.** Heatmaps representing the co-occurrence of phyla in sister groups to LECA clades for **A)** informational processing and **B)** metabolic gene trees. Co-occurrences were considered at topological distance 3, when the taxonomic proportion of query phylum is higher than 25%. Numbers represent the sum of taxonomic proportions for the respective co-occurences. Only those sums of taxonomic proportions of co-occurrences that sum more than 2 are represented in the heatmap. Extended data for **Fig. 1C,D.**

**Figure S3.** Specific co-occurrences in informational processing gene trees involving eukaryotic parasites. Co-occurrences of **A)** Dependentiae and Asgardarchaeota and **B)** Depedentiae and Chlamydiota.

**Figure S4.** Presence of prokaryotic phyla in sister groups to LECA clades for metabolic subcategories denoted by KEGG: **A)** amino acid metabolism, **B)** carbohydrate metabolism, **C)** energy metabolism, **D)** lipid metabolism, **E)** metabolism of cofactors and vitamins. The quantification combines cumulative topological distances 1, 2, 3 when taxonomic proportion is higher than 50% and ultrafast bootstrap support equal or higher than 70%, which represent an approximation of actual contributions.

**Figure S5.** Pruned phylogenetic trees representing potential archaeal origins related with amino acid metabolisms. Tip labels of collapsed branches indicate the most abundant presence of taxa. In the same order as in labels: bacteria or archaea, total number of sequences (also indicated between brackets), number of archaeal (A-), bacterial (B-) and asgardarchaeal (Asg-) sequences, the most abundant prokaryotic phyla, and the proportion of most abundant prokaryotic phyla. For eukaryotic tip labels: KO ids, KO gene name, number of Excavata (E-), Amorphea (A-) and Diaphoretickes (D-) sequences, and total number of sequences between brackets. Only those sister groups that are at topological distance 6 to the LECA node are shown.

**Figure S6.** Pruned phylogenetic trees representing potential archaeal origins related to carbohydrate metabolism. See above for tip label explanation.

**Figure S7.** Pruned phylogenetic trees representing potential archaeal origins related to lipid metabolism. See above for tip label explanation.

**Figure S8.** Pruned phylogenetic trees representing potential archaeal origins related to metabolism of cofactors and vitamins. See above for tip label explanation.

**Figure S9.** Pruned phylogenetic trees representing potential archaeal origins related to nucleotide metabolism. See above for tip label explanation.

**Figure S10.** Pruned phylogenetic trees representing potential archaeal origins related to glycan metabolism. See above for tip label explanation.

**Figure S11.** Taxonomic contributions for diverse functional categories related to other metabolism gene families. Only the top 6 phyla were shown for each category. The sisterhood relationship was considered when taxonomic proportion is higher than 50% and ultrafast bootstrap support equal or higher than 70. Cumulative topological distances 1, 2 and 3, were shown independently.

**Figure S12.** Taxonomic contributions for diverse functional categories related with other informational processing gene families. Only the top 6 phyla were shown for each category. The sisterhood relationship was considered when taxonomic proportion is higher than 50% and ultrafast bootstrap support equal or higher than 70. Cumulative topological distances 1, 2 and 3, were shown independently.

**Figure S13.** Relative timing of potential gene contributions during eukaryogenesis using normalized stem branch-length, different size of LECA (>= 10 or >=60 eukaryotic taxa) clades and different number of taxa allowed in the sister group (>=1 or >1). Upper panels show the distribution of stem branch-lengths for all gene trees, mid panel for informational processing related gene trees, and lower panel for metabolism related gene trees. Density plot, and box plot show the same data for the stem branch-length distribution, while barplots represent the number of sisterhood relationships for the respective stem-branchlength distribution. Only sisterhood relationships at topological distance

1, involving more than 1 sequence, with taxonomic proportion higher than 70% and ultrafast bootstrap than 70%, were analyzed.

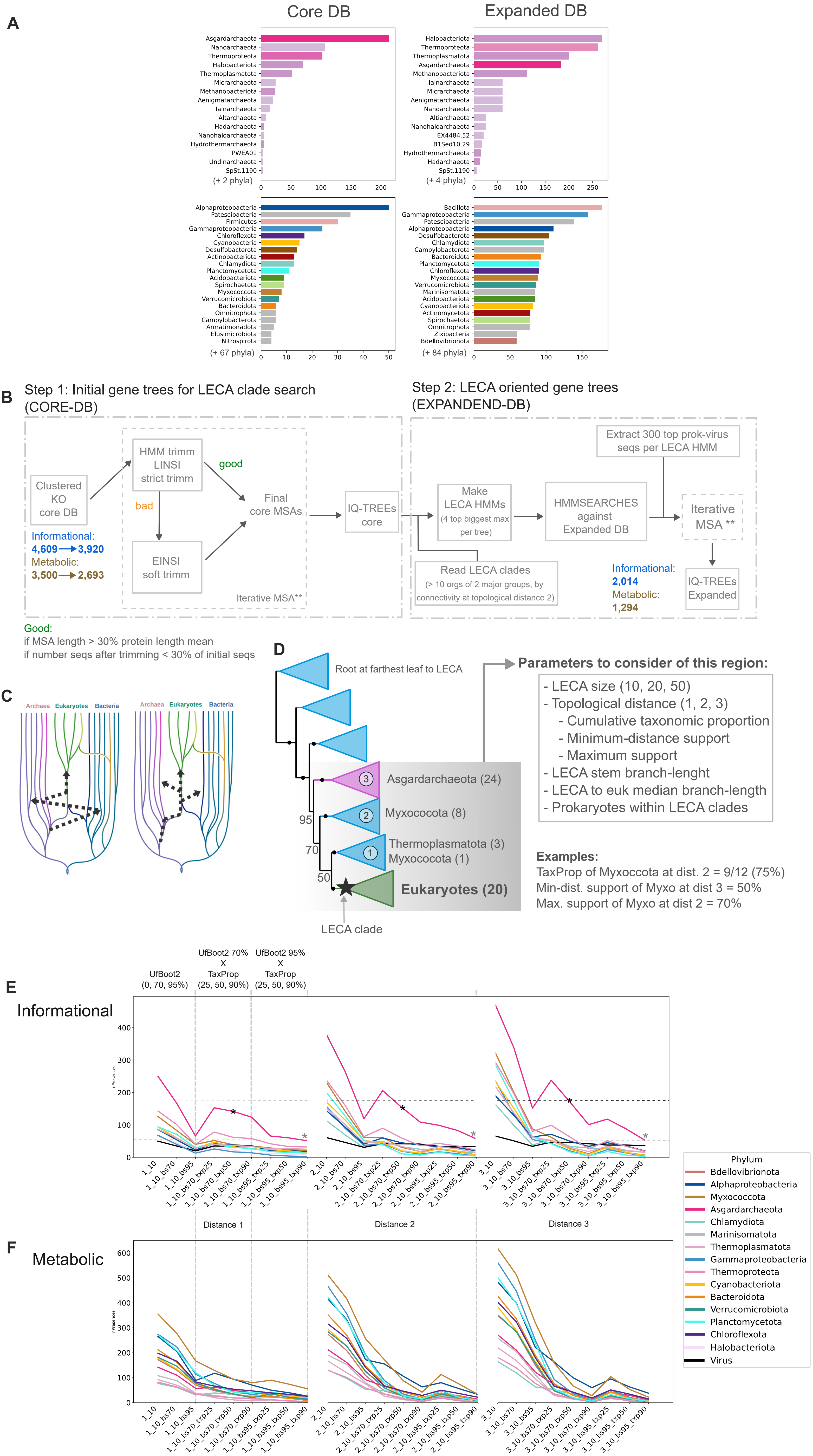
A
Core DB
Expanded DB
Asgardarchaeota
Nanoarchaeota
Thermoproteota
Halobacteriota
Thermoplasmatota
Micrarchaeota
Methanobacteriota
Aenigmatarchaeota
Iainarchaeota
Altarchaeota
Hadarchaeota
Nanohaloarchaeota
Hydrothermarchaeota
PWEA01
Undinarchaeota
SpSt.1190
(+ 2 phyla)
(+ 4 phyla)
(+ 67 phyla)
(+ 84 phyla)
B
Step 1: Initial gene trees for LECA clade search (CORE-DB)
Step 2: LECA oriented gene trees (EXPANDEND-DB)
Clustered KO core DB
Informational:
4,609 → 3,920
Metabolic:
3,500 → 2,693
HMM trimm LINSI strict trimm
good
bad
EINSI soft trimm
Final core MSAs
Iterative MSA**
IQ-TREEs core
Make LECA HMMs (4 top biggest max per tree)
Read LECA clades (> 10 orgs of 2 major groups, by connectivity at topological distance 2)
HMMSEARCHES against Expanded DB
Extract 300 top prok-virus seqs per LECA HMM
Iterative MSA **
IQ-TREEs Expanded
Informational:
2,014
Metabolic:
1,294
Good:
if MSA length > 30% protein length mean
if number seqs after trimming < 30% of initial seqs
C
Archaea
Eukaryotes
Bacteria
D
Root at farthest leaf to LECA
Asgardarchaeota (24)
Myxococota (8)
Thermoplasmatota (3)
Myxococota (1)
Eukaryotes (20)
95
70
50
LECA clade
Parameters to consider of this region:
- LECA size (10, 20, 50)
- Topological distance (1, 2, 3)
- Cumulative taxonomic proportion
- Minimum-distance support
- Maximum support
- LECA stem branch-lenght
- LECA to euk median branch-length
- Prokaryotes within LECA clades
Examples:
TaxProp of Myxoccota at dist. 2 = 9/12 (75%)
Min-dist. support of Myxo at dist 3 = 50%
Max. support of Myxo at dist 2 = 70%
E
Informational
UfBoot2 (0, 70, 95%)
UfBoot2 70% X TaxProp (25, 50, 90%)
UfBoot2 95% X TaxProp (25, 50, 90%)
nPresences
Distance 1
Distance 2
Distance 3
F
Metabolic
Method
Phylum
Bdellovibrionota
Alphaproteobacteria
Myxococcota
Asgardarchaeota
Chlamydiota
Marinisomatota
Thermoplasmatota
Gammaproteobacteria
Thermoproteota
Cyanobacteriota
Bacteroidota
Verrucomicrobiota
Planctomycetota
Chloroflexota
Halobacteriota
Virus

Fig. S1

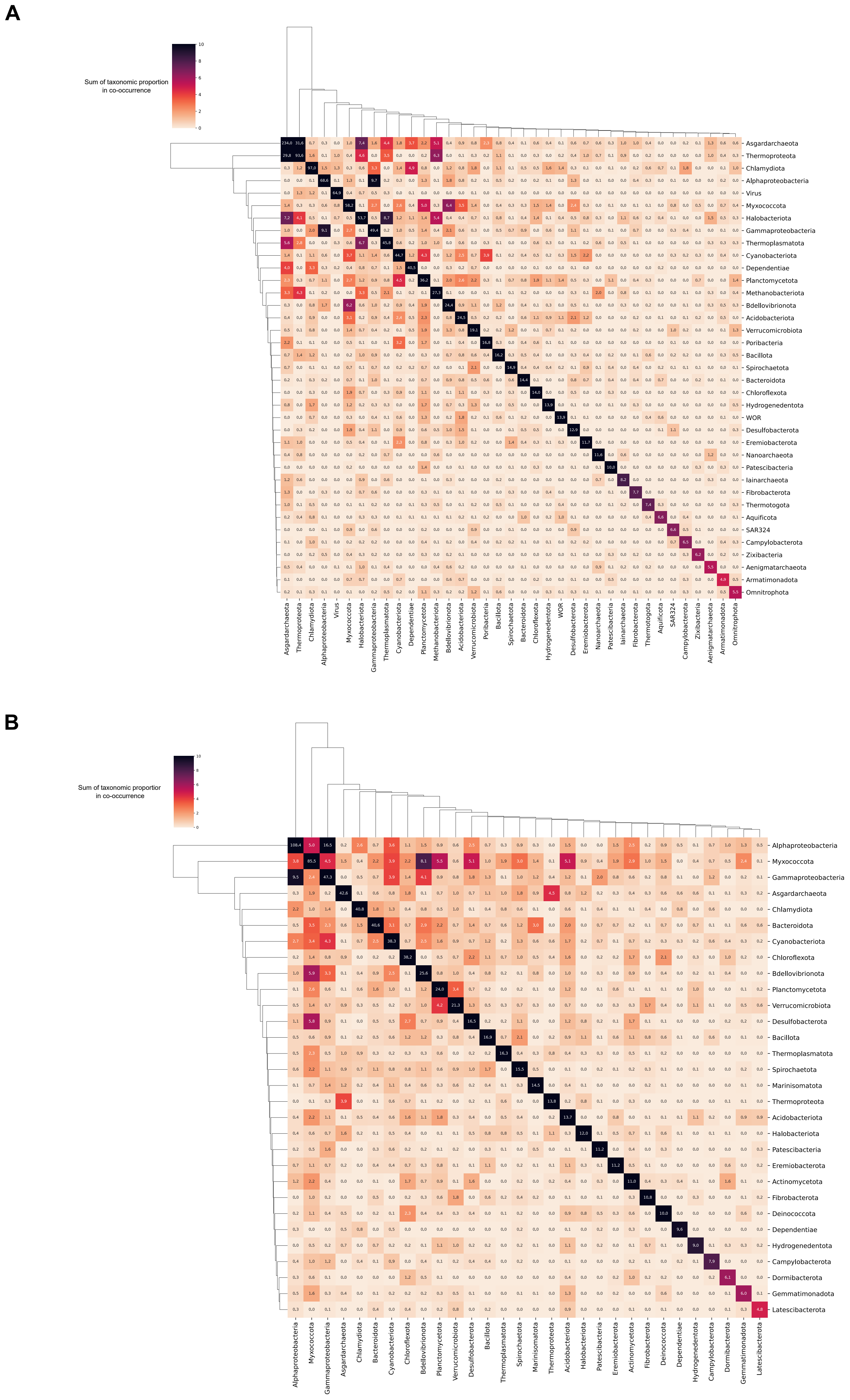
A
Sum of taxonomic proportion
in co-occurrence
0
2
4
6
8
10
Asgardarchaeota
Thermoproteota
Chlamydiota
Alphaproteobacteria
Virus
Myxococcota
Halobacteriota
Gammaproteobacteria
Thermoplasmatota
Cyanobacteriota
Dependentiae
Planctomycetota
Methanobacteriota
Bdellovibrionota
Acidobacteriota
Verrucomicrobiota
Poribacteria
Bacillota
Spirochaetota
Bacteroidota
Chloroflexota
Hydrogenedentota
WOR
Desulfobacterota
Eremiobacterota
Nanoarchaeota
Patescibacteria
Iainarchaeota
Fibrobacterota
Thermotogota
Aquificota
SAR324
Campylobacterota
Zixibacteria
Aenigmatarchaeota
Armatimonadota
Omnitrophota
B
Sum of taxonomic proportior
in co-occurrence
0
2
4
6
8
10
Alphaproteobacteria
Myxococcota
Gammaproteobacteria
Asgardarchaeota
Chlamydiota
Bacteroidota
Cyanobacteriota
Chloroflexota
Bdellovibrionota
Planctomycetota
Verrucomicrobiota
Desulfobacterota
Bacillota
Thermoplasmatota
Spirochaetota
Marinisomatota
Thermoproteota
Acidobacteriota
Halobacteriota
Patescibacteria
Eremiobacterota
Actinomycetota
Fibrobacterota
Deinococcota
Dependentiae
Hydrogenedentota
Campylobacterota
Dormibacterota
Gemmatimonadota
Latescibacterota

Fig. S2

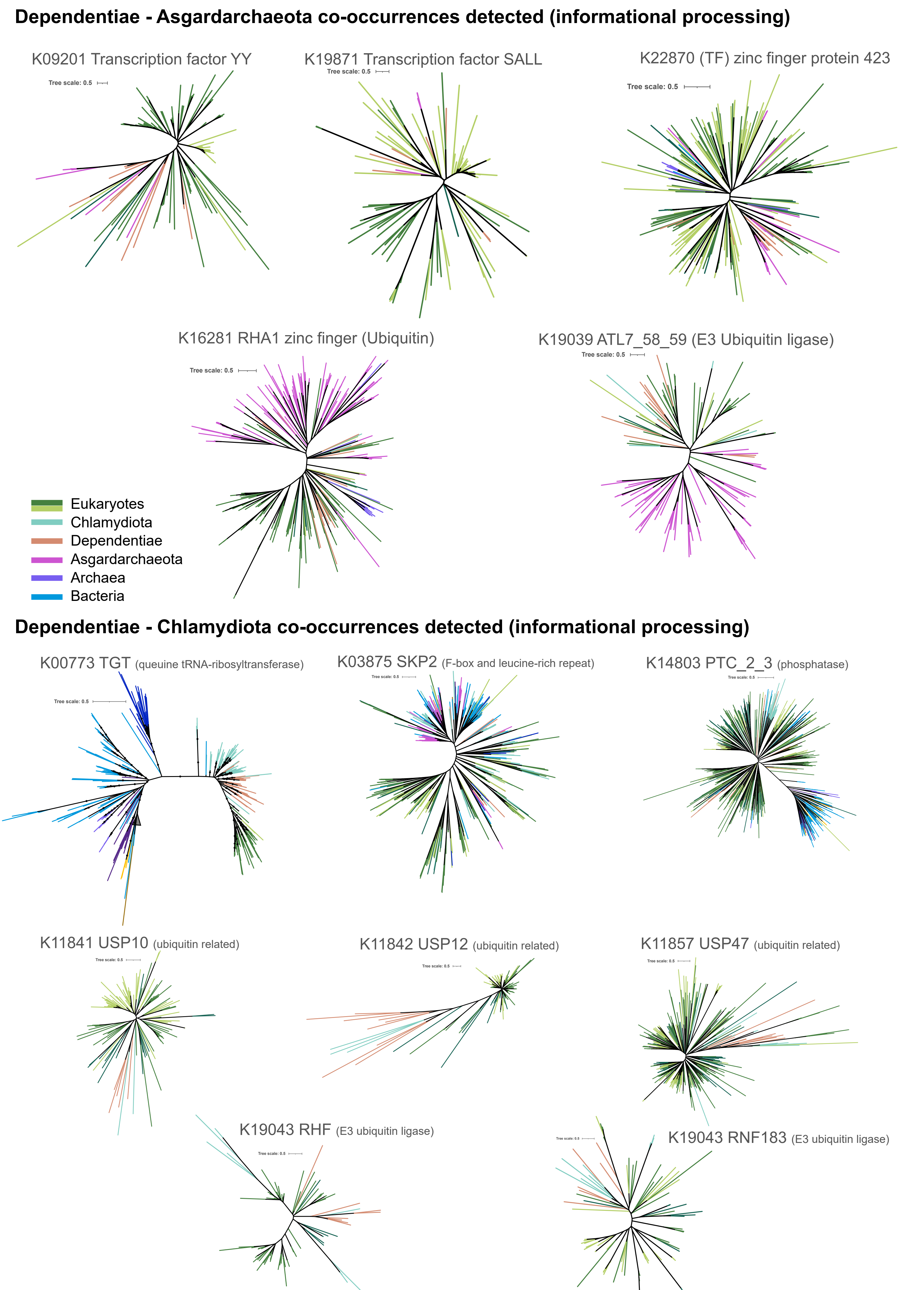
Dependentiae - Asgardarchaeota co-occurrences detected (informational processing)
K09201 Transcription factor YY
Tree scale: 0.5
K19871 Transcription factor SALL
Tree scale: 0.5
K22870 (TF) zinc finger protein 423
Tree scale: 0.5
K16281 RHA1 zinc finger (Ubiquitin)
Tree scale: 0.5
K19039 ATL7_58_59 (E3 Ubiquitin ligase)
Tree scale: 0.5
Eukaryotes
Chlamydiota
Dependentiae
Asgardarchaeota
Archaea
Bacteria
Dependentiae - Chlamydiota co-occurrences detected (informational processing)
K00773 TGT (queuine tRNA-ribosyltransferase)
Tree scale: 0.5
K03875 SKP2 (F-box and leucine-rich repeat)
K14803 PTC_2_3 (phosphatase)
K11841 USP10 (ubiquitin related)
K11842 USP12 (ubiquitin related)
K11857 USP47 (ubiquitin related)
K19043 RHF (E3 ubiquitin ligase)
K19043 RNF183 (E3 ubiquitin ligase)

Fig. S3

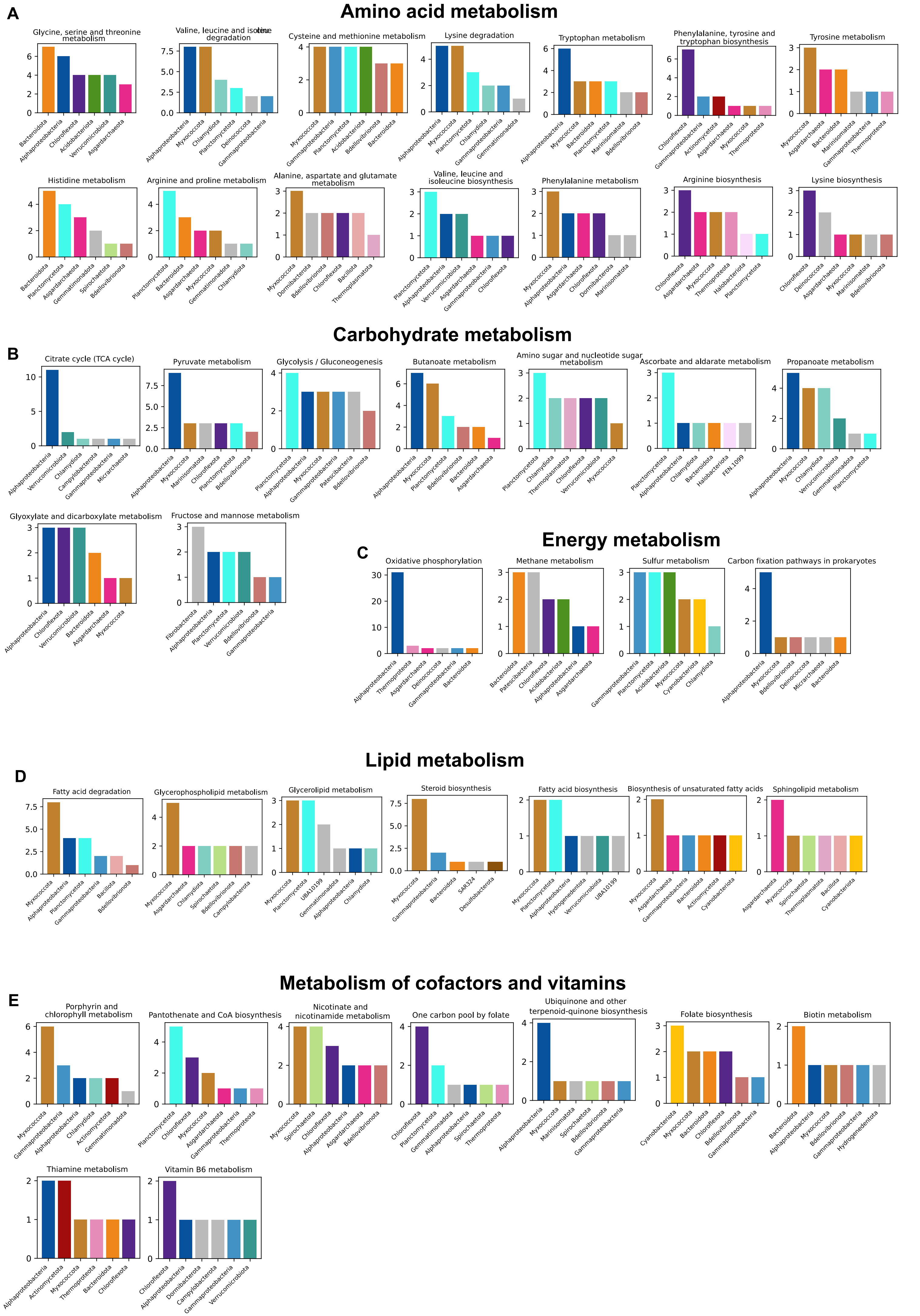
A
Amino acid metabolism
Glycine, serine and threonine metabolism
Bacteroidota, Alphaproteobacteria, Chloroflexota, Acidobacteriota, Verrucomicrobiota, Asgardarchaeota
Valine, leucine and isoleucine degradation
Alphaproteobacteria, Myxococcota, Chlamydiota, Planctomycetota, Deinococcota, Gammaproteobacteria
Cysteine and methionine metabolism
Myxococcota, Gammaproteobacteria, Planctomycetota, Acidobacteriota, Bdellovibrionota, Bacteroidota
Lysine degradation
Alphaproteobacteria, Myxococcota, Planctomycetota, Chlamydiota, Gammaproteobacteria, Gemmatimonadota
Tryptophan metabolism
Alphaproteobacteria, Myxococcota, Bacteroidota, Planctomycetota, Marinisomatota, Bdellovibrionota
Phenylalanine, tyrosine and tryptophan biosynthesis
Chloroflexota, Gammaproteobacteria, Actinomycetota, Asgardarchaeota, Myxococcota, Thermoproteota
Tyrosine metabolism
Myxococcota, Asgardarchaeota, Bacteroidota, Marinisomatota, Gammaproteobacteria, Thermoproteota
Histidine metabolism
Bacteroidota, Planctomycetota, Asgardarchaeota, Gemmatimonadota, Spirochaetota, Bdellovibrionota
Arginine and proline metabolism
Planctomycetota, Bacteroidota, Asgardarchaeota, Myxococcota, Gemmatimonadota, Chlamydiota
Alanine, aspartate and glutamate metabolism
Myxococcota, Dormibacterota, Bdellovibrionota, Chloroflexota, Bacillota, Thermoplasmatota
Valine, leucine and isoleucine biosynthesis
Planctomycetota, Alphaproteobacteria, Verrucomicrobiota, Asgardarchaeota, Gammaproteobacteria, Chloroflexota
Phenylalanine metabolism
Myxococcota, Alphaproteobacteria, Asgardarchaeota, Chloroflexota, Dormibacterota, Marinisomatota
Arginine biosynthesis
Chloroflexota, Asgardarchaeota, Myxococcota, Thermoproteota, Halobacteriota, Planctomycetota
Lysine biosynthesis
Chloroflexota, Deinococcota, Asgardarchaeota, Myxococcota, Marinisomatota, Bdellovibrionota
Carbohydrate metabolism
B
Citrate cycle (TCA cycle)
Alphaproteobacteria, Verrucomicrobiota, Chlamydiota, Campylobacterota, Gammaproteobacteria, Micrarchaeota
Pyruvate metabolism
Alphaproteobacteria, Myxococcota, Marinisomatota, Chloroflexota, Planctomycetota, Bdellovibrionota
Glycolysis / Gluconeogenesis
Planctomycetota, Alphaproteobacteria, Myxococcota, Gammaproteobacteria, Patescibacteria, Bdellovibrionota
Butanoate metabolism
Alphaproteobacteria, Myxococcota, Planctomycetota, Bdellovibrionota, Bacteroidota, Asgardarchaeota
Amino sugar and nucleotide sugar metabolism
Planctomycetota, Chlamydiota, Thermoplasmatota, Chloroflexota, Verrucomicrobiota, Myxococcota
Ascorbate and aldarate metabolism
Planctomycetota, Alphaproteobacteria, Chlamydiota, Bacteroidota, Halobacteriota, FEN-1099
Propanoate metabolism
Alphaproteobacteria, Myxococcota, Chlamydiota, Verrucomicrobiota, Gemmatimonadota, Planctomycetota
Glyoxylate and dicarboxylate metabolism
Alphaproteobacteria, Chloroflexota, Verrucomicrobiota, Bacteroidota, Asgardarchaeota, Myxococcota
Fructose and mannose metabolism
Fibrobacterota, Alphaproteobacteria, Planctomycetota, Verrucomicrobiota, Bdellovibrionota, Gammaproteobacteria
Energy metabolism
C
Oxidative phosphorylation
Alphaproteobacteria, Thermoproteota, Asgardarchaeota, Deinococcota, Gammaproteobacteria, Bacteroidota
Methane metabolism
Bacteroidota, Patescibacteria, Chloroflexota, Acidobacteriota, Alphaproteobacteria, Asgardarchaeota
Sulfur metabolism
Gammaproteobacteria, Planctomycetota, Acidobacteriota, Myxococcota, Cyanobacteriota, Chlamydiota
Carbon fixation pathways in prokaryotes
Alphaproteobacteria, Myxococcota, Bdellovibrionota, Deinococcota, Micrarchaeota, Bacteroidota
Lipid metabolism
D
Fatty acid degradation
Myxococcota, Alphaproteobacteria, Planctomycetota, Gammaproteobacteria, Bacillota, Bdellovibrionota
Glycerophospholipid metabolism
Myxococcota, Asgardarchaeota, Chlamydiota, Spirochaetota, Bdellovibrionota, Campylobacterota
Glycerolipid metabolism
Myxococcota, Planctomycetota, UBA10199, Gemmatimonadota, Alphaproteobacteria, Chlamydiota
Steroid biosynthesis
Myxococcota, Gammaproteobacteria, Bacteroidota, SAR324, Desulfobacterota
Fatty acid biosynthesis
Myxococcota, Planctomycetota, Alphaproteobacteria, Hydrogenedentota, Verrucomicrobiota, UBA10199
Biosynthesis of unsaturated fatty acids
Myxococcota, Asgardarchaeota, Gammaproteobacteria, Bacteroidota, Actinomycetota, Cyanobacteriota
Sphingolipid metabolism
Asgardarchaeota, Myxococcota, Spirochaetota, Thermoplasmatota, Bacillota, Cyanobacteriota
Metabolism of cofactors and vitamins
E
Porphyrin and chlorophyll metabolism
Myxococcota, Gammaproteobacteria, Alphaproteobacteria, Chlamydiota, Actinomycetota, Gemmatimonadota
Pantothenate and CoA biosynthesis
Planctomycetota, Chloroflexota, Myxococcota, Asgardarchaeota, Gammaproteobacteria, Thermoproteota
Nicotinate and nicotinamide metabolism
Myxococcota, Spirochaetota, Chloroflexota, Alphaproteobacteria, Asgardarchaeota, Bdellovibrionota
One carbon pool by folate
Chloroflexota, Planctomycetota, Gemmatimonadota, Alphaproteobacteria, Spirochaetota, Thermoproteota
Ubiquinone and other terpenoid-quinone biosynthesis
Alphaproteobacteria, Myxococcota, Marinisomatota, Spirochaetota, Bdellovibrionota, Gammaproteobacteria
Folate biosynthesis
Cyanobacteriota, Myxococcota, Bacteroidota, Chloroflexota, Bdellovibrionota, Gammaproteobacteria
Biotin metabolism
Bacteroidota, Alphaproteobacteria, Myxococcota, Bdellovibrionota, Gammaproteobacteria, Hydrogenedentota
Thiamine metabolism
Alphaproteobacteria, Actinomycetota, Myxococcota, Thermoproteota, Bacteroidota, Chloroflexota
Vitamin B6 metabolism
Chloroflexota, Alphaproteobacteria, Dormibacterota, Campylobacterota, Gammaproteobacteria, Verrucomicrobiota

Fig. S4

### Archaeal origin of amino acid metabolism

#### Glutamate family

**Arginine:**
OTC, K00611
argE, K01438
arcA, K01478

**Lysine:**
LYS9, K00293

#### Aromatic family

**Phenylalanine:**
paaD, K02612

Phe-Tyr-Tryp:
hisC, K00817

#### Histidine

HutL, K01468

MULTIFUNCTIONAL
**Cys-Met., Val-Leu-Isoleu,**
**Gly-Threo-Ser:**
SDS, K17989
**Gly-Ser-Threo:**
TDH, K15789
**Arg-Pro, Ala, Hist:**
CNDP2, K08660

OTHERS
**Selenocompound:**
SEPSECS, K03341

**Glutathione:**
ggt, K00681

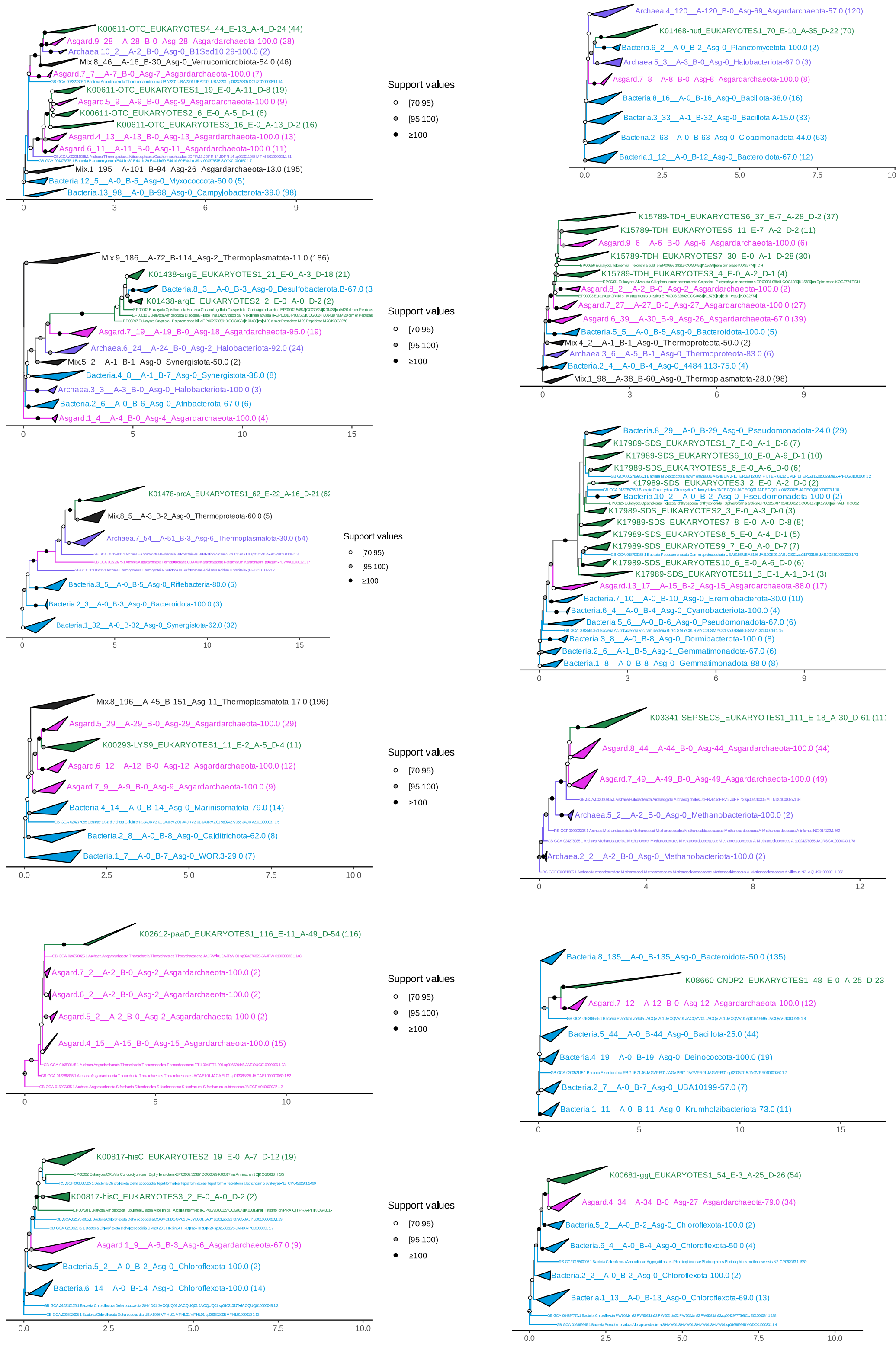


**Fig. S5**

**Glycolysis**
GpmI, K15633
apgM, K15635
ENO, K01689
ADPGK, K08074

**Amino-sugar /nucleotide-sugar**
GMPP, K00966

**Glyoxylate/ dicarboxylate**
PGP, K19269

**Inositol phosphate**
ITPK1, K00913

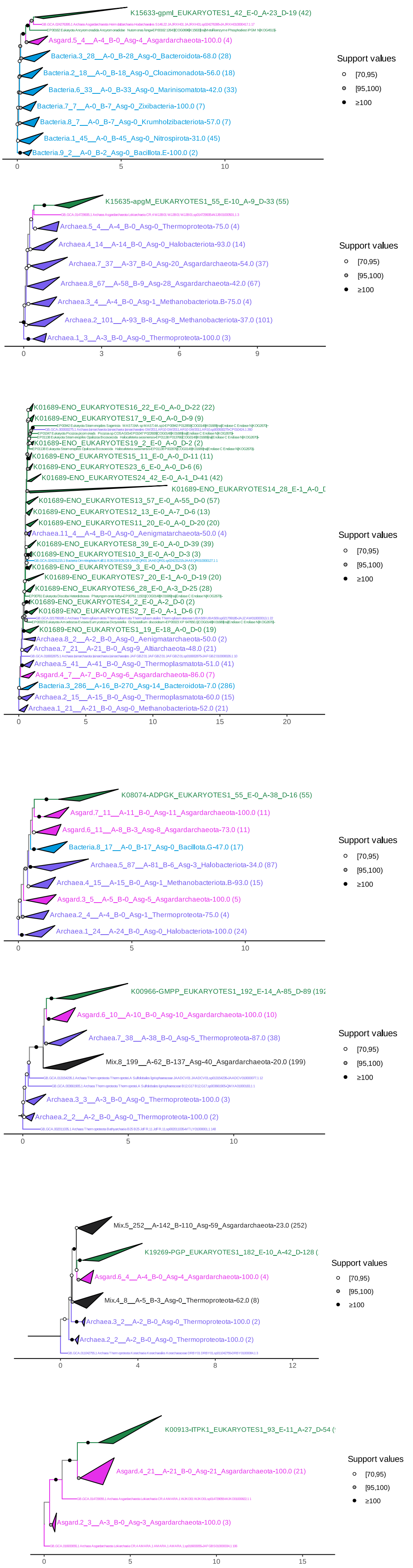


**Fig. S6**

Archaeal origins of Lipid metabolism

#### Glycerophospholipid

PLMT, K00550
GPD1, K00006

#### Sphingolipid

GBA2, K17108
SGPL1, K01634
SGPP1, K04716

#### Unsat. fatty acid

TesA, K10804

#### Mevalonate-Isoprenoid

HMGS, K01641
MVK, K00869

*Nested in Archaea:*
MVAK2, K00938
MVD, K01597
IPK, K06981
GGPS1, K00804
DHDDS, K11778
HMGR, K00021

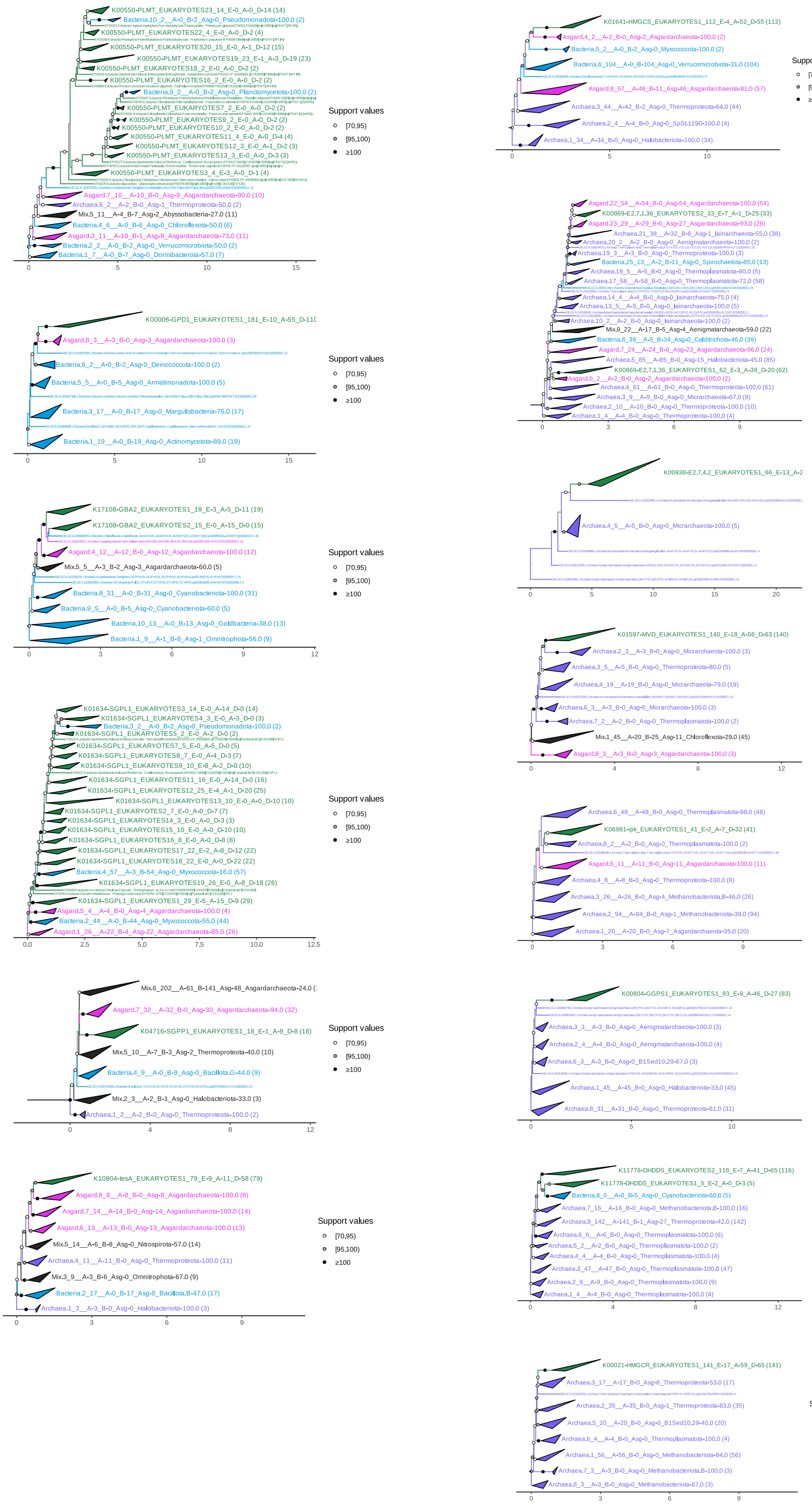


Fig. S7

### Archaeal origin of metabolism of cofactors and vitamins

**Nicotinate/Nicotinamide**
nadC, K00767

**One carbon pool folate**
MTFHS, K01934

**Pantothenate and CoA**
PANK_1_2_3, K09680
E2.7.7.3B, K02201
COASY, K02318

**Thiamine metabolism**
THI4, K03146

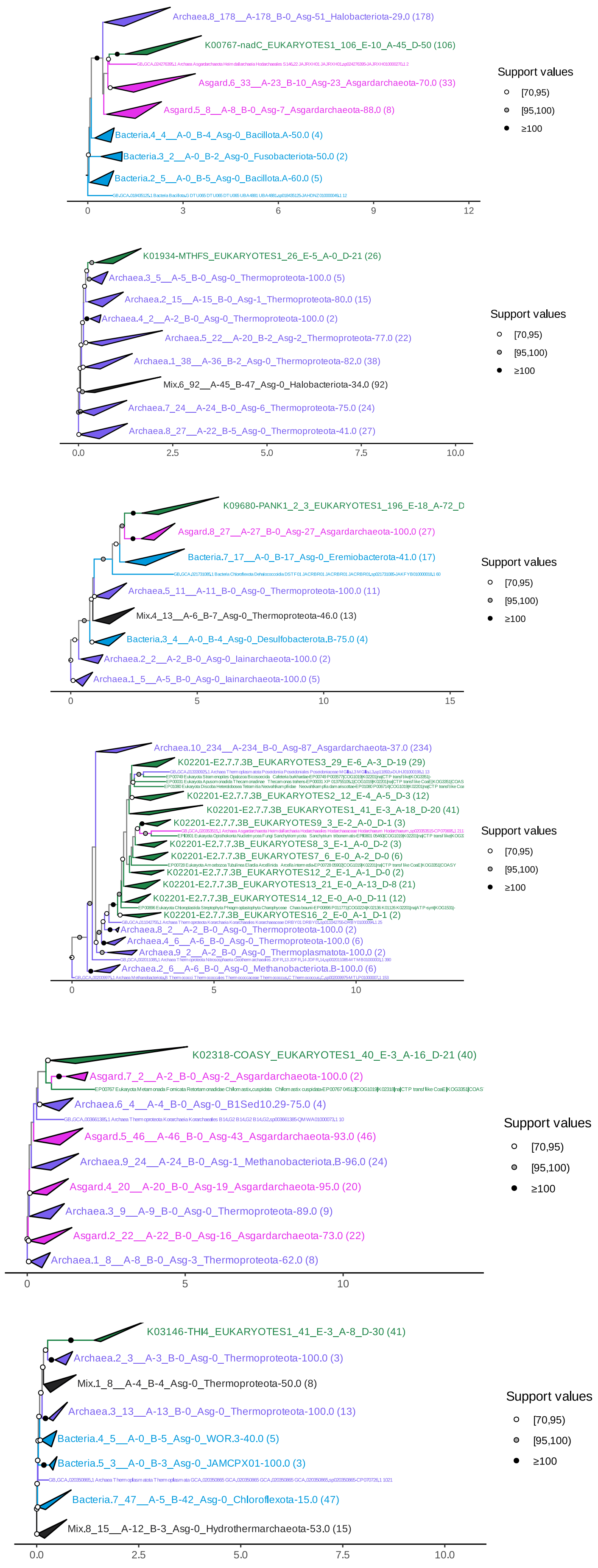


**Fig. S8**

**Purine-Pyrimidine**
rtpR, K00527

**Pyrimidine**
*Nested in Archaea:*
tmk, K00943
pyrG, K01937
udp, K00757 * root issue

**Purine**
AK6, K18532

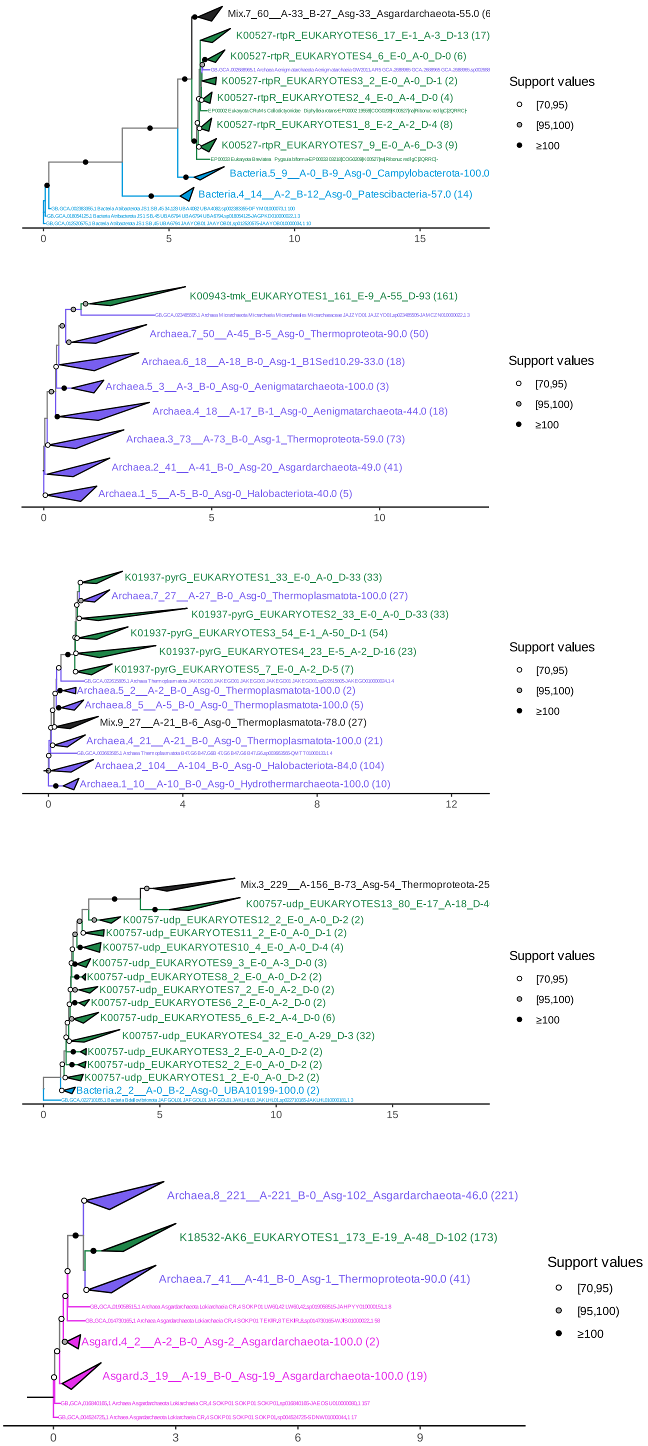


**Fig. S9**

**GPI-anchor biosynthesis**
PIGA, K03857
PIGB, K05286
PIGO, K05288
DPM2, K09658

**N-Glycan biosynthesis**
ALG13, K07432
ALG14, K07441
ALG11, K03844
ALG2, K03843
ALG7, K01001

OST1, K12666
STT3, K07151
OST3, K12669

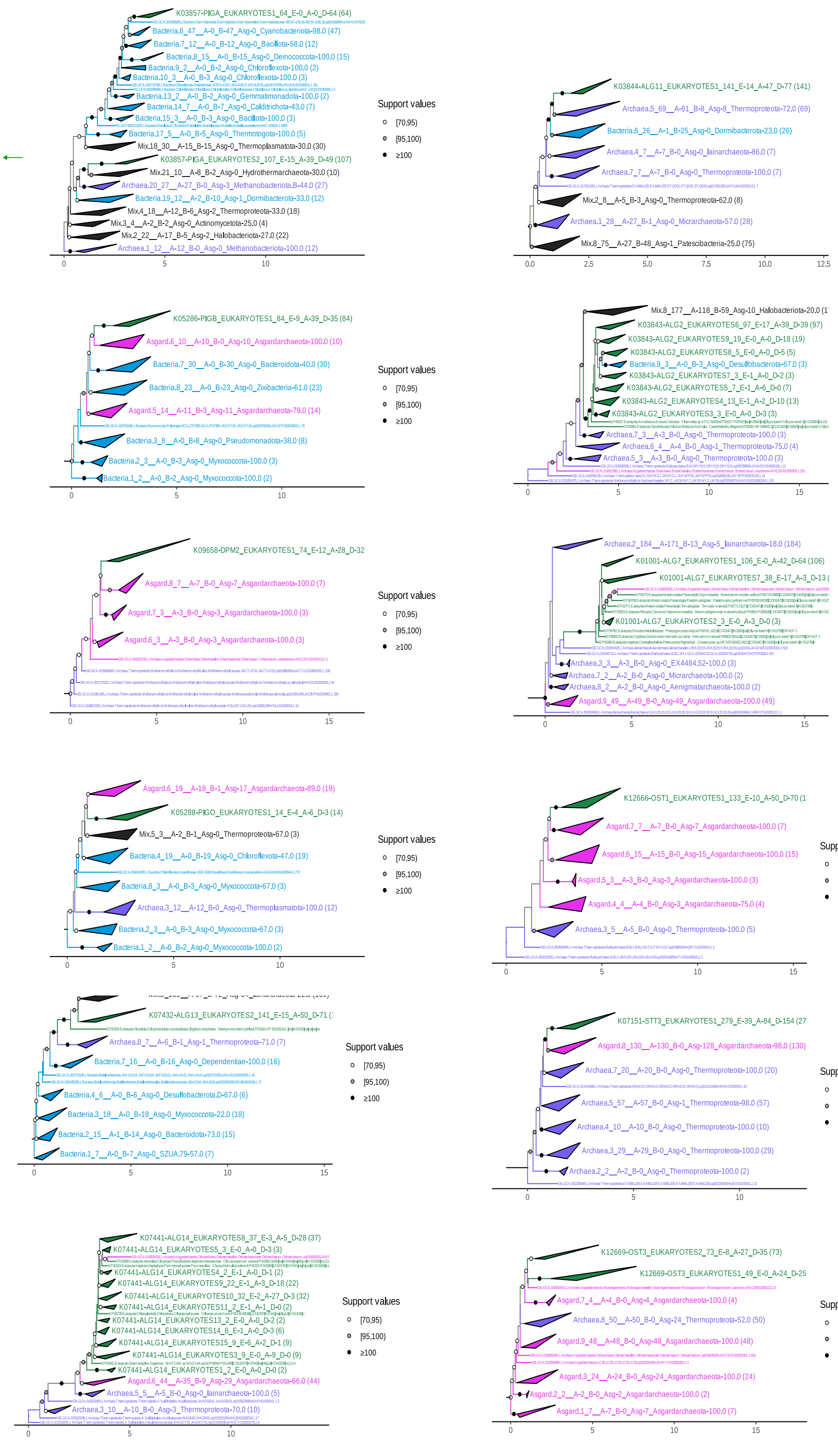


**Fig. S10**

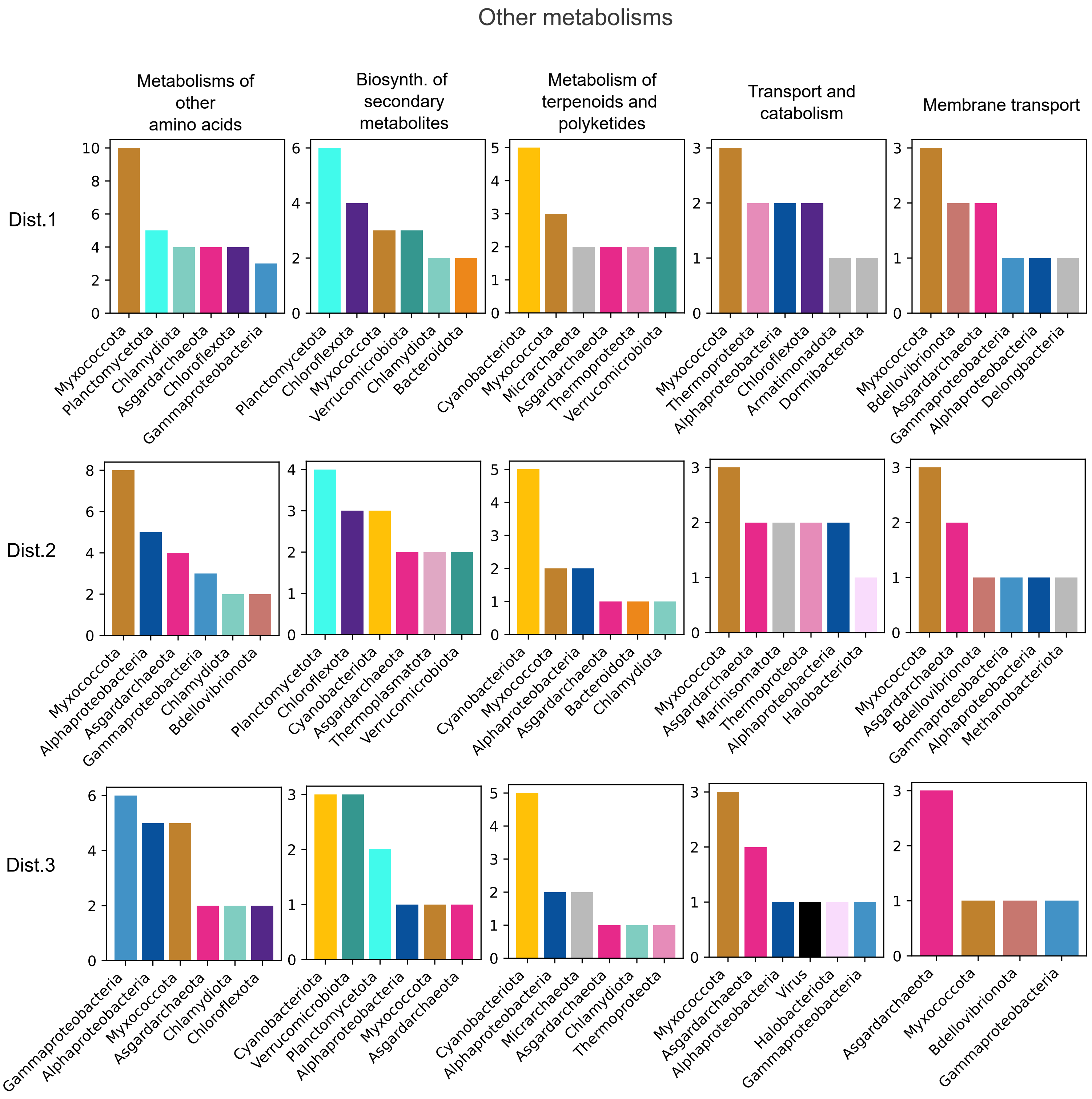
Other metabolisms
Metabolisms of other amino acids
Biosynth. of secondary metabolites
Metabolism of terpenoids and polyketides
Transport and catabolism
Membrane transport
Dist.1
Dist.2
Dist.3
Myxococcota
Planctomycetota
Chlamydiota
Asgardarchaeota
Chloroflexota
Gammaproteobacteria
Verrucomicrobiota
Bacteroidota
Cyanobacteriota
Micrarchaeota
Thermoproteota
Alphaproteobacteria
Armatimonadota
Dormibacterota
Bdellovibrionota
Delongbacteria
Thermoplasmatota
Marinisomatota
Halobacteriota
Methanobacteriota
Virus

**Fig. S11**

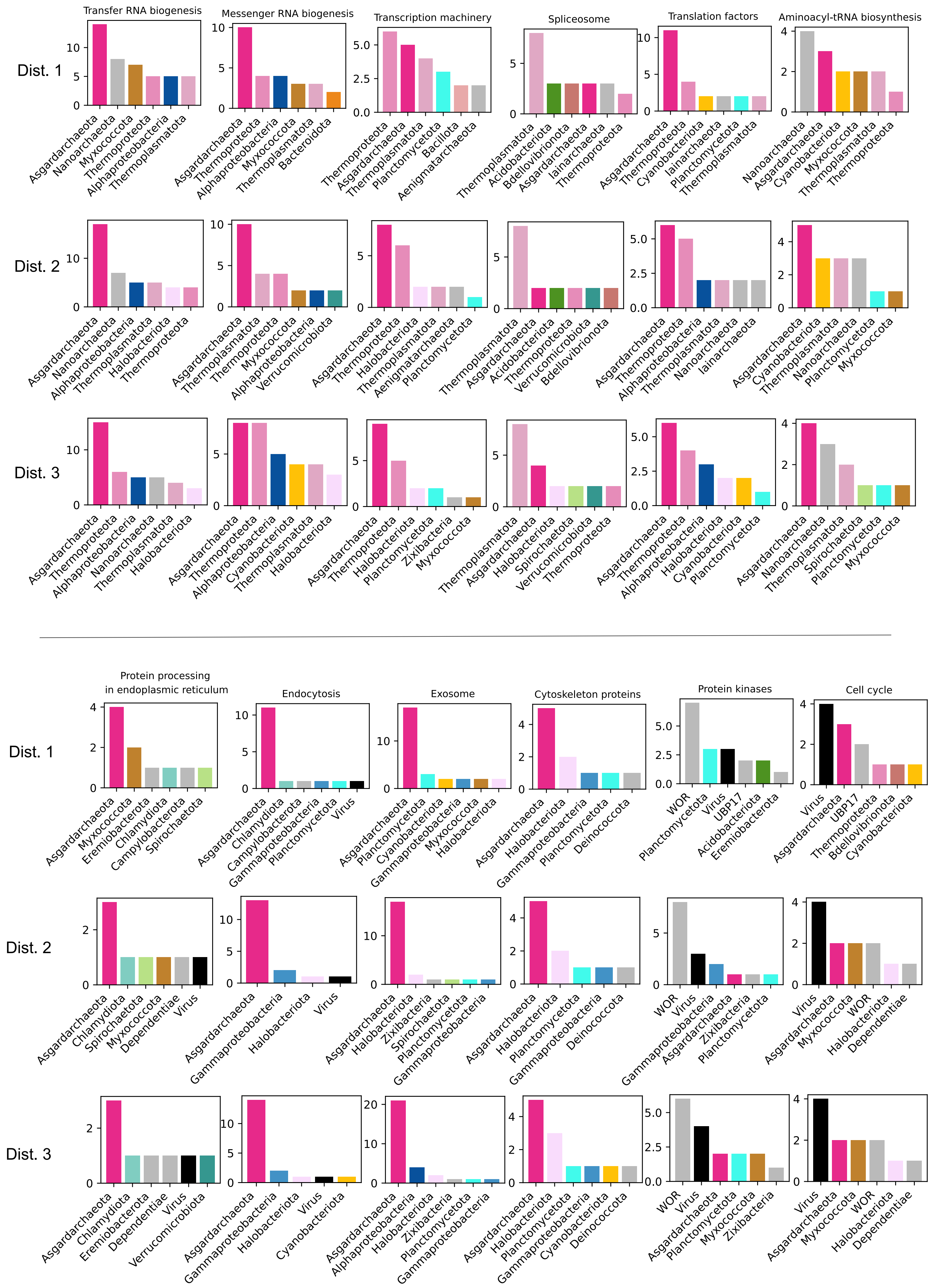
Transfer RNA biogenesis
Messenger RNA biogenesis
Transcription machinery
Spliceosome
Translation factors
Aminoacyl-tRNA biosynthesis
Dist. 1
Asgardarchaeota
Nanoarchaeota
Myxococcota
Thermoproteota
Alphaproteobacteria
Thermoplasmatota
Asgardarchaeota
Thermoproteota
Alphaproteobacteria
Myxococcota
Thermoplasmatota
Bacteroidota
Thermoproteota
Asgardarchaeota
Thermoplasmatota
Planctomycetota
Bacillota
Aenigmatarchaeota
Thermoplasmatota
Acidobacteriota
Bdellovibrionota
Asgardarchaeota
Iainarchaeota
Thermoproteota
Asgardarchaeota
Thermoproteota
Cyanobacteriota
Iainarchaeota
Planctomycetota
Thermoplasmatota
Nanoarchaeota
Asgardarchaeota
Cyanobacteriota
Myxococcota
Thermoplasmatota
Thermoproteota
Dist. 2
Asgardarchaeota
Nanoarchaeota
Alphaproteobacteria
Thermoplasmatota
Halobacteriota
Thermoproteota
Asgardarchaeota
Thermoplasmatota
Thermoproteota
Myxococcota
Alphaproteobacteria
Verrucomicrobiota
Asgardarchaeota
Thermoproteota
Halobacteriota
Thermoplasmatota
Aenigmatarchaeota
Planctomycetota
Thermoplasmatota
Asgardarchaeota
Acidobacteriota
Thermoproteota
Verrucomicrobiota
Bdellovibrionota
Asgardarchaeota
Thermoproteota
Alphaproteobacteria
Thermoplasmatota
Nanoarchaeota
Iainarchaeota
Asgardarchaeota
Cyanobacteriota
Thermoplasmatota
Nanoarchaeota
Planctomycetota
Myxococcota
Dist. 3
Asgardarchaeota
Thermoproteota
Alphaproteobacteria
Nanoarchaeota
Thermoplasmatota
Halobacteriota
Asgardarchaeota
Thermoproteota
Alphaproteobacteria
Cyanobacteriota
Thermoplasmatota
Halobacteriota
Asgardarchaeota
Thermoproteota
Halobacteriota
Planctomycetota
Zixibacteria
Myxococcota
Thermoplasmatota
Asgardarchaeota
Halobacteriota
Spirochaetota
Verrucomicrobiota
Thermoproteota
Asgardarchaeota
Thermoproteota
Alphaproteobacteria
Halobacteriota
Cyanobacteriota
Planctomycetota
Asgardarchaeota
Nanoarchaeota
Thermoplasmatota
Spirochaetota
Planctomycetota
Myxococcota
Protein processing
in endoplasmic reticulum
Endocytosis
Exosome
Cytoskeleton proteins
Protein kinases
Cell cycle
Dist. 1
Asgardarchaeota
Myxococcota
Eremiobacterota
Chlamydiota
Campylobacterota
Spirochaetota
Asgardarchaeota
Chlamydiota
Campylobacterota
Gammaproteobacteria
Planctomycetota
Virus
Asgardarchaeota
Planctomycetota
Cyanobacteriota
Gammaproteobacteria
Myxococcota
Halobacteriota
Asgardarchaeota
Halobacteriota
Gammaproteobacteria
Planctomycetota
Deinococcota
WOR
Planctomycetota
Virus
UBP17
Acidobacteriota
Eremiobacterota
Virus
Asgardarchaeota
UBP17
Thermoproteota
Bdellovibrionota
Cyanobacteriota
Dist. 2
Asgardarchaeota
Chlamydiota
Spirochaetota
Myxococcota
Dependentiae
Virus
Asgardarchaeota
Gammaproteobacteria
Halobacteriota
Virus
Asgardarchaeota
Halobacteriota
Zixibacteria
Spirochaetota
Planctomycetota
Gammaproteobacteria
Asgardarchaeota
Halobacteriota
Planctomycetota
Gammaproteobacteria
Deinococcota
WOR
Virus
Gammaproteobacteria
Asgardarchaeota
Zixibacteria
Planctomycetota
Virus
Asgardarchaeota
Myxococcota
WOR
Halobacteriota
Dependentiae
Dist. 3
Asgardarchaeota
Chlamydiota
Eremiobacterota
Dependentiae
Virus
Verrucomicrobiota
Asgardarchaeota
Gammaproteobacteria
Halobacteriota
Virus
Cyanobacteriota
Asgardarchaeota
Alphaproteobacteria
Halobacteriota
Zixibacteria
Planctomycetota
Gammaproteobacteria
Asgardarchaeota
Halobacteriota
Planctomycetota
Gammaproteobacteria
Cyanobacteriota
Deinococcota
WOR
Virus
Asgardarchaeota
Planctomycetota
Myxococcota
Zixibacteria
Virus
Asgardarchaeota
Myxococcota
WOR
Halobacteriota
Dependentiae

**Fig. S12**

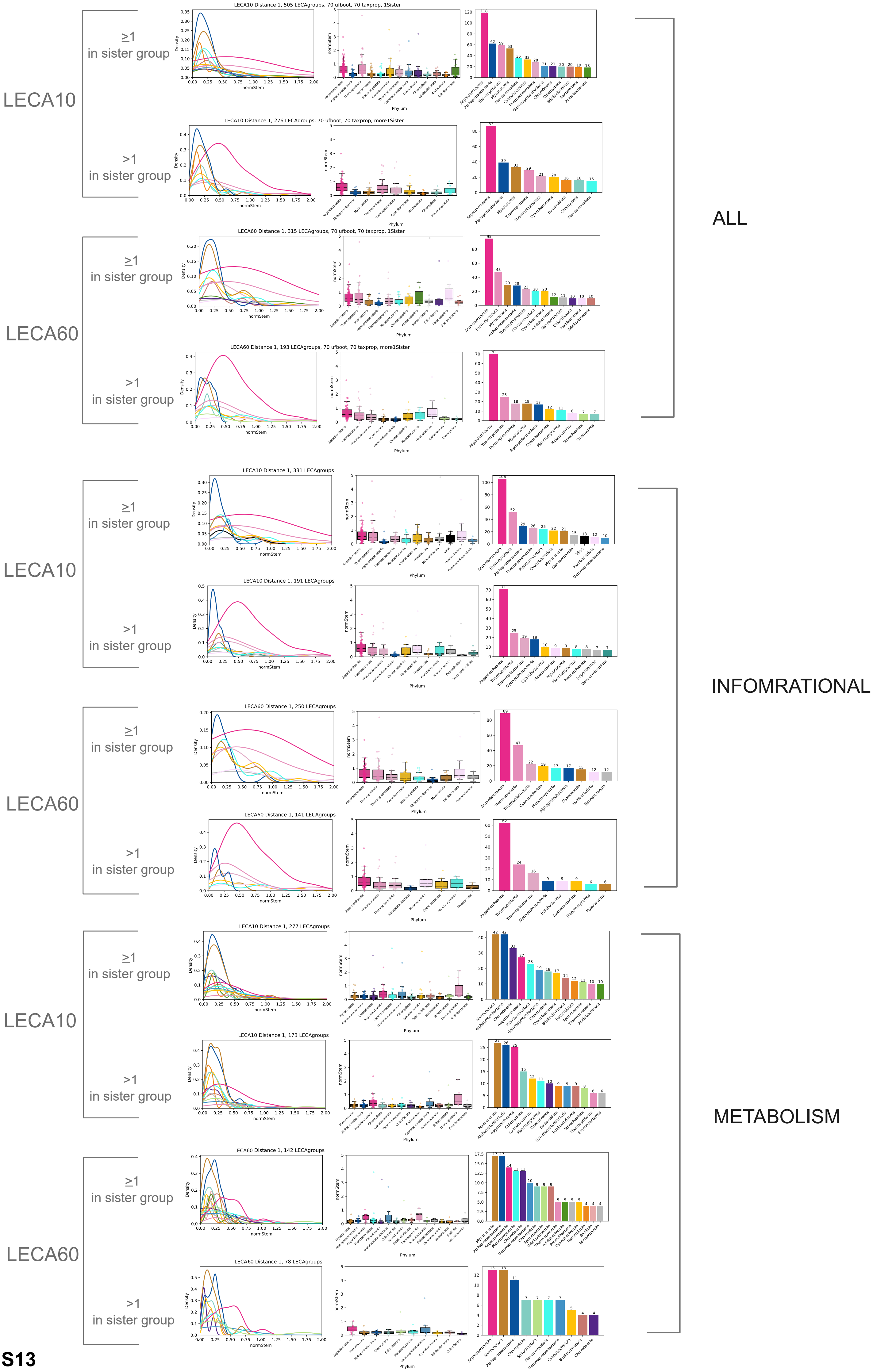

LECA10
LECA60
≥1 in sister group
>1 in sister group
ALL
INFOMRATIONAL
METABOLISM
LECA10 Distance 1, 505 LECAgroups, 70 ufboot, 70 taxprop, 1Sister
LECA10 Distance 1, 276 LECAgroups, 70 ufboot, 70 taxprop, more1Sister
LECA60 Distance 1, 315 LECAgroups, 70 ufboot, 70 taxprop, 1Sister
LECA60 Distance 1, 193 LECAgroups, 70 ufboot, 70 taxprop, more1Sister
LECA10 Distance 1, 331 LECAgroups
LECA10 Distance 1, 191 LECAgroups
LECA60 Distance 1, 250 LECAgroups
LECA60 Distance 1, 141 LECAgroups
LECA10 Distance 1, 277 LECAgroups
LECA10 Distance 1, 173 LECAgroups
LECA60 Distance 1, 142 LECAgroups
LECA60 Distance 1, 78 LECAgroups
Density
normStem
Phylum


Fig. S13
